# Comparative profiling of carnitine palmitoyltransferase 1 isoforms reveals vincamine as a selective carnitine palmitoyltransferase 1b inhibitor

**DOI:** 10.64898/2026.05.23.727424

**Authors:** Anthony Wong, William Luo, Justin Xuan, Himanshu Gupta, Matthew Li, Anusha Natraj, Saharsh Madullapalli, Helen Tao, Connor Wahng, Maanav Balan, Melinda Wu, Zhan Chen

## Abstract

Carnitine palmitoyltransferase 1 (CPT1) catalyzes the rate-limiting step of fatty acid oxidation and has emerged as a therapeutic target for metabolic diseases and cancer. CPT1 exists in three isoforms, CPT1a, CPT1b, and CPT1c, with distinct tissue distributions and enzymatic properties; however, limitations of previous platforms enabling parallel isoform comparison have undermined efforts to identify selective inhibitors that could minimize off-target effects. Here, we describe a DTNB-based enzyme activity assay adapted for high-throughput screening of CPT1b, the predominant isoform in cardiac and skeletal muscle. Mitochondrial extracts from Expi293F cells transfected with CPT1a or CPT1b expression plasmids served as sources of catalytically active enzymes. The assay was validated using three previously confirmed CPT1b inhibitors: (R)-(+)-etomoxir, perhexiline, and malonyl-CoA. We then generated side-by-side inhibitory profiles for both isoforms, identifying vincamine as a lead selective inhibitor of CPT1b. Furthermore, chlorpromazine, previously characterized only as a broad CPT1 inhibitor and subsequently shown to inhibit CPT1a, is demonstrated here to also inhibit CPT1b, expanding its known isoform profile. Together, these results establish a robust platform for comparative isoform profiling and demonstrate that selective modulation of CPT1b is achievable, with implications for targeted therapeutics in metabolic and oncological disease.

## Introduction

Fatty acid oxidation (FAO) is a key metabolic process that breaks long-chain fatty acids into acetyl-CoA, which enters the tricarboxylic acid (TCA) cycle to drive ATP production (1). Before β-oxidation can occur, free fatty acids (FFAs) are transported into the cell via long-chain acyl-CoA synthetase, forming long-chain acyl-CoA (2). Because acyl-CoA cannot cross the inner mitochondrial membrane, carnitine palmitoyltransferase 1 (CPT1) converts acyl-CoA into acyl-carnitine for mitochondrial import (2). CPT1 comprises three isoforms (CPT1a and CPT1b on the outer mitochondrial membrane, with CPT1c in the endoplasmic reticulum), with CPT1a and CPT1b catalyzing the rate-limiting step (3). CPT1b predominates in tissues with high FAO demand in the muscle and heart, specializing in fatty acid β-oxidation and accounting for ∼98% of cardiac CPT1 activity and ∼95% of skeletal muscle CPT1 activity (4).

Inhibiting CPT1b effectively blocks FAO in muscle and cardiac tissue (5). This shifts cellular metabolism toward greater reliance on glycolysis, which may limit ATP production during periods of high energy demand (4). The altered metabolic state impacts overall energy balance, changes substrate utilization, and may lead to lipid accumulation upstream (6). In addition, impaired FAO can cause FGF21 expression in skeletal muscle, which increases glucose uptake (7).

Therapeutically, partial suppression of FAO via CPT1 inhibition can shift substrate use toward glucose oxidation, allowing for treatment of obesity (8) and type 2 diabetes (9) by reducing lipid overload and improving insulin sensitivity (8). CPT1 activity also plays a relevant role in cancer: CPT1 inhibition sensitizes cancerous cells to apoptosis (10, 11, 12).

Given the critical role of CPT1 in energy metabolism, it is essential to distinguish among its isoforms. CPT1 isoforms exhibit different enzymatic kinetics and inhibitory sensitivity to small-molecule inhibitors (13). Because CPT1a and CPT1b share a 63% amino acid identity and have similar catalytic roles, comparative analyses between them provide significant insight into isoform-specific regulation of FAO (3). Specifically, CPT1a exhibits a higher affinity for carnitine [Km = 30 µM in rat CPT1a] compared to CPT1b [Km = 500 µM in rat CPT1b] (3). However, CPT1b demonstrates approximately 100-fold greater sensitivity to malonyl-CoA inhibition when compared to CPT1a (5). These distinctions likely reflect tissue-specific adaptations to metabolic demand and other functions.

Understanding these distinctions is crucial for clarifying tissue-specific roles and pharmacological relevance, as non-selective CPT1 inhibitors can disrupt essential metabolic functions across multiple tissues, leading to severe off-target side effects such as cardiac hypertrophy, seizures, oxidative stress (14), and death (15, 16). The biological significance of a CPT1a-CPT1b comparative assay can be attributed to the isoforms’ distinct physiological roles. While CPT1a has been extensively studied in hepatic and cancer metabolism (17), comparatively minimal work has examined the regulatory function and inhibitory sensitivity of CPT1b in direct comparison to CPT1a. Given the tissue-specific localization of CPT1b (4), the lack of isoform-specific investigation limits both mechanistic insight and therapeutic development (18). To address this gap, we used CPT1a and CPT1b enzymes expressed in Expi293F cells to screen for isoform-selective compounds under biologically relevant conditions, enabling direct evaluation of differential inhibition between the two isoforms.

## Results

### CPT1b Expression, Purification, and Quantification using Immunoassay

ELISA was used to quantify the protein expression level of CPT1b in transfected Expi293F cells and compared against non-transfected controls. CPT1b was expressed at a higher level than CPT1a. Together, the ELISA, Bradford, and TOM20 results were used to normalize expression levels across CPT1a and CPT1b samples, allowing a more accurate and precise determination of isoform selectivity in subsequent enzymatic assays (Figure 1).

**Figure 1.**
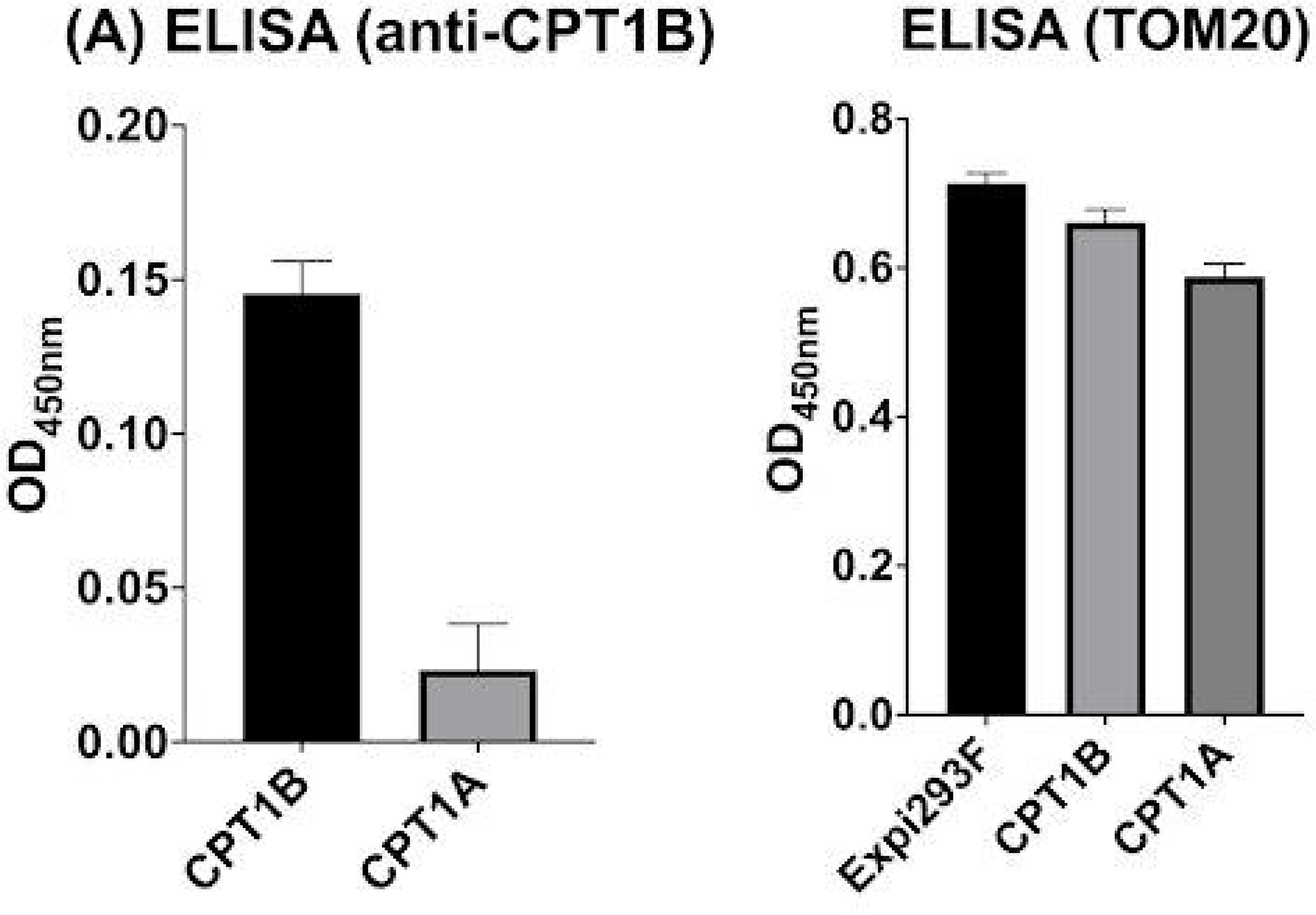
Quantification of CPT1 isoforms in transgenic Expi293F cells. ELISA analysis shows higher mitochondrial CPT1b protein levels compared to CPT1a (**A**). TOM20 showed comparable protein levels between Expi293F, CPT1b, and CPT1a samples. Data are represented as mean ± SEM (n=3), and significance was calculated using a Welch’s t-test (** p < 0.01 and **** p < 0.0001).

### Validating CPT1b Enzyme Activity and Inhibitor Sensitivity Assay

Etomoxir was used to ensure the DTNB-modified platform is reliable in comparing CPT1a and CPT1b enzyme activity and the inhibitor sensitivity assay (Figure 2) (19). Additionally, DMSO alone had no detectable effect on CPT1b activity, confirming that the observed inhibition is compound-specific (Figure 5).

**Figure 2.**
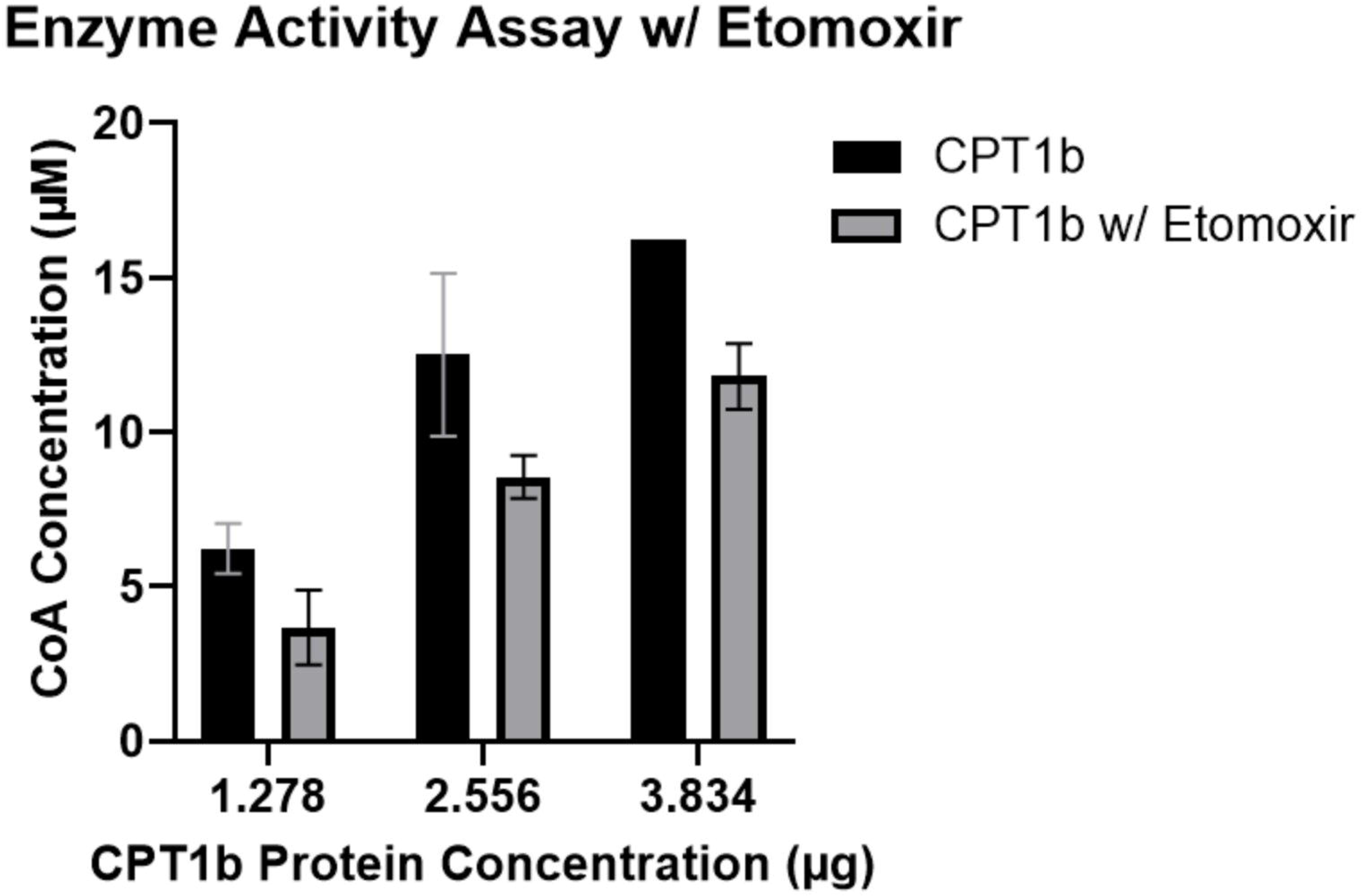
CPT1b activity assay and (R)-(+)-etomoxir inhibitor sensitivity assay. The same cell lysis pellet solution was used with a protein concentration of 12.78 μg/μL with varying concentrations of 2.56, 5.12, and 7.68 μg/μL. 10 μL of a 10 mM (R)-(+)-etomoxir stock solution was added to each group to achieve a final concentration of 100 µM. Data are represented as mean (n=3) and significance was calculated using a Welch’s t-test (** p < 0.01 and **** p < 0.0001).

A series of previously reported CPT1 inhibitors, including (R)-(+)-etomoxir (20), malonyl-CoA (21), and perhexiline (22), were used to test the modified DTNB-based spectrophotometric assay utilizing mitochondrial extracts from CPT1b-transfected Expi293F cells [Figure 3].

**Figure 3.**
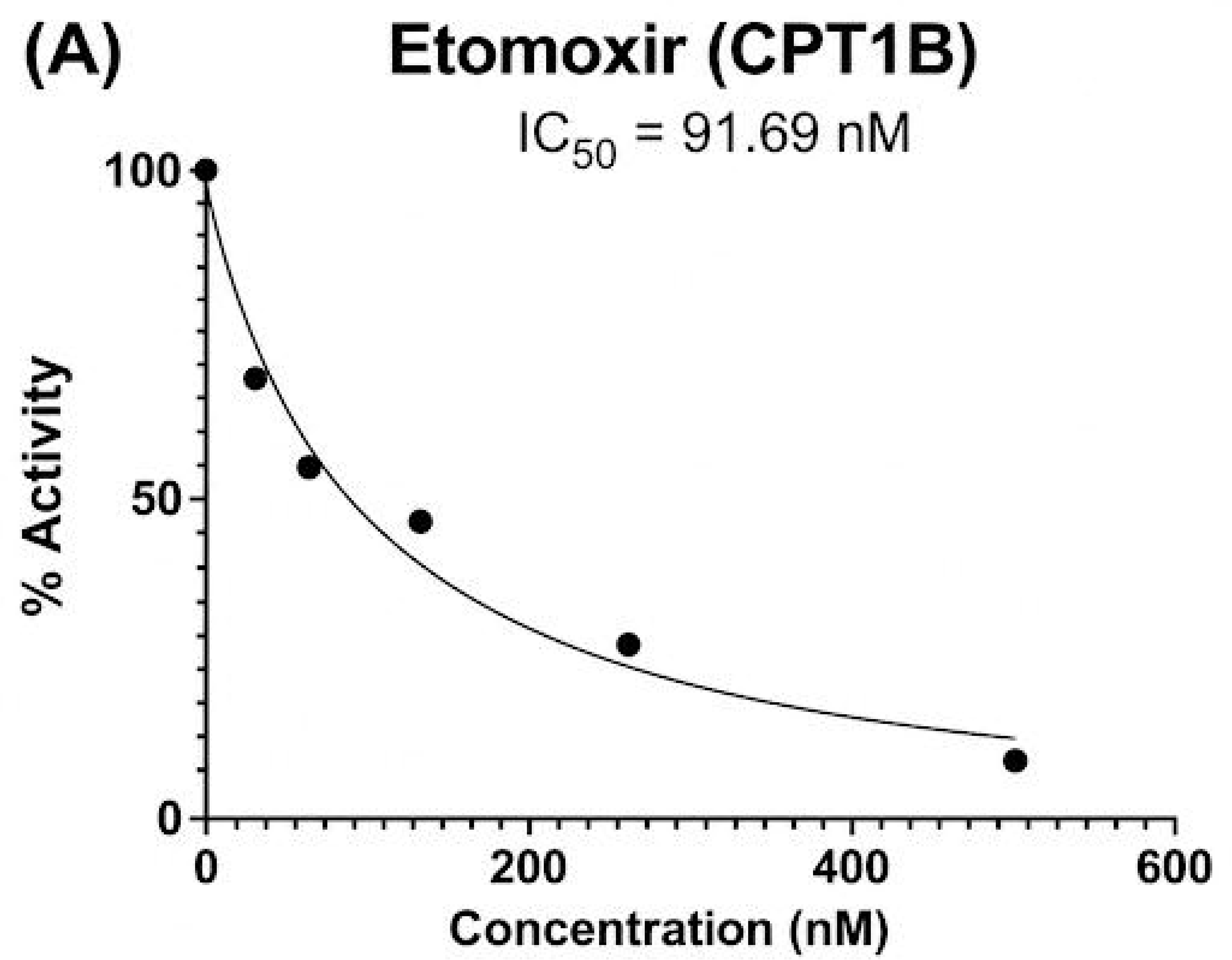
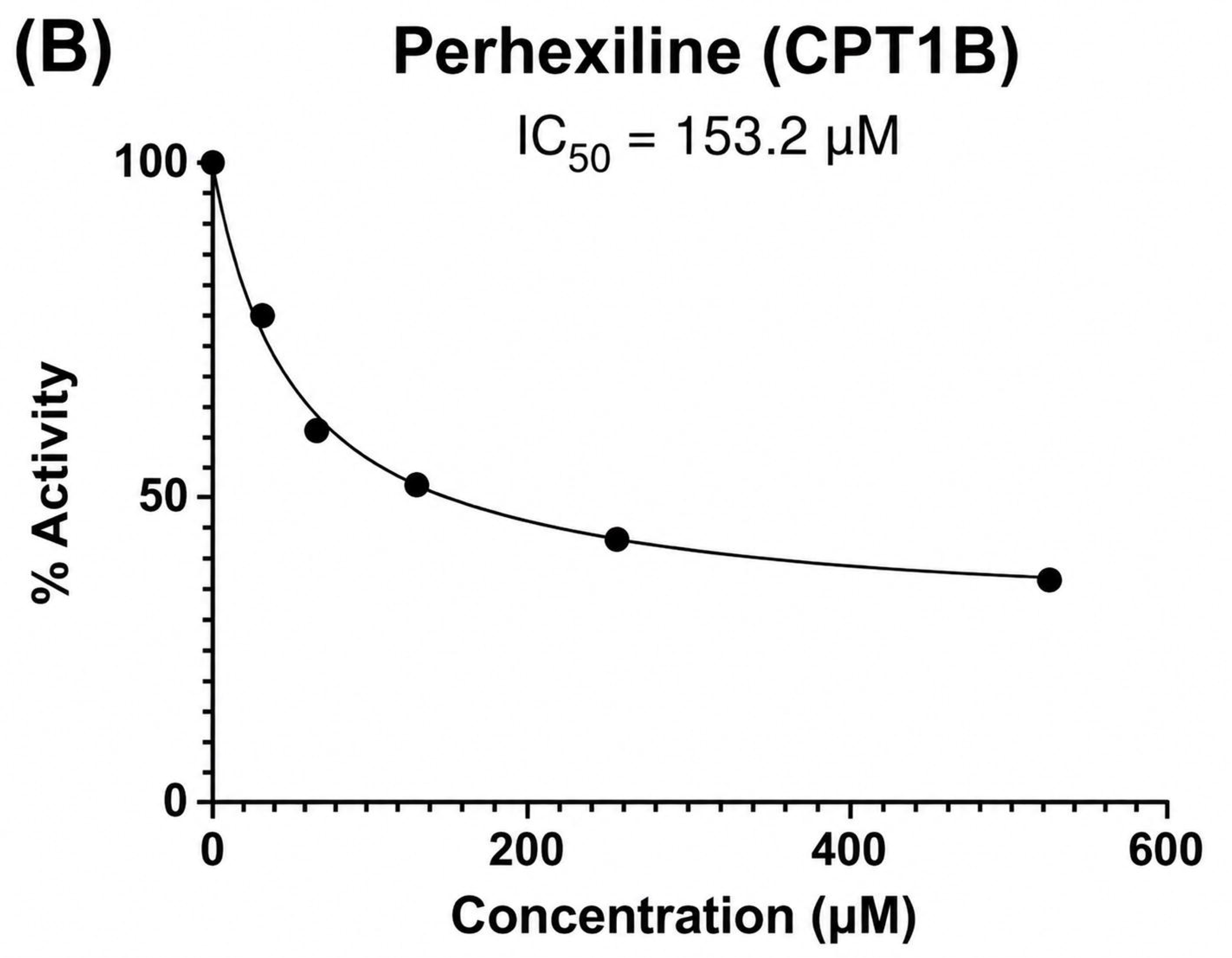
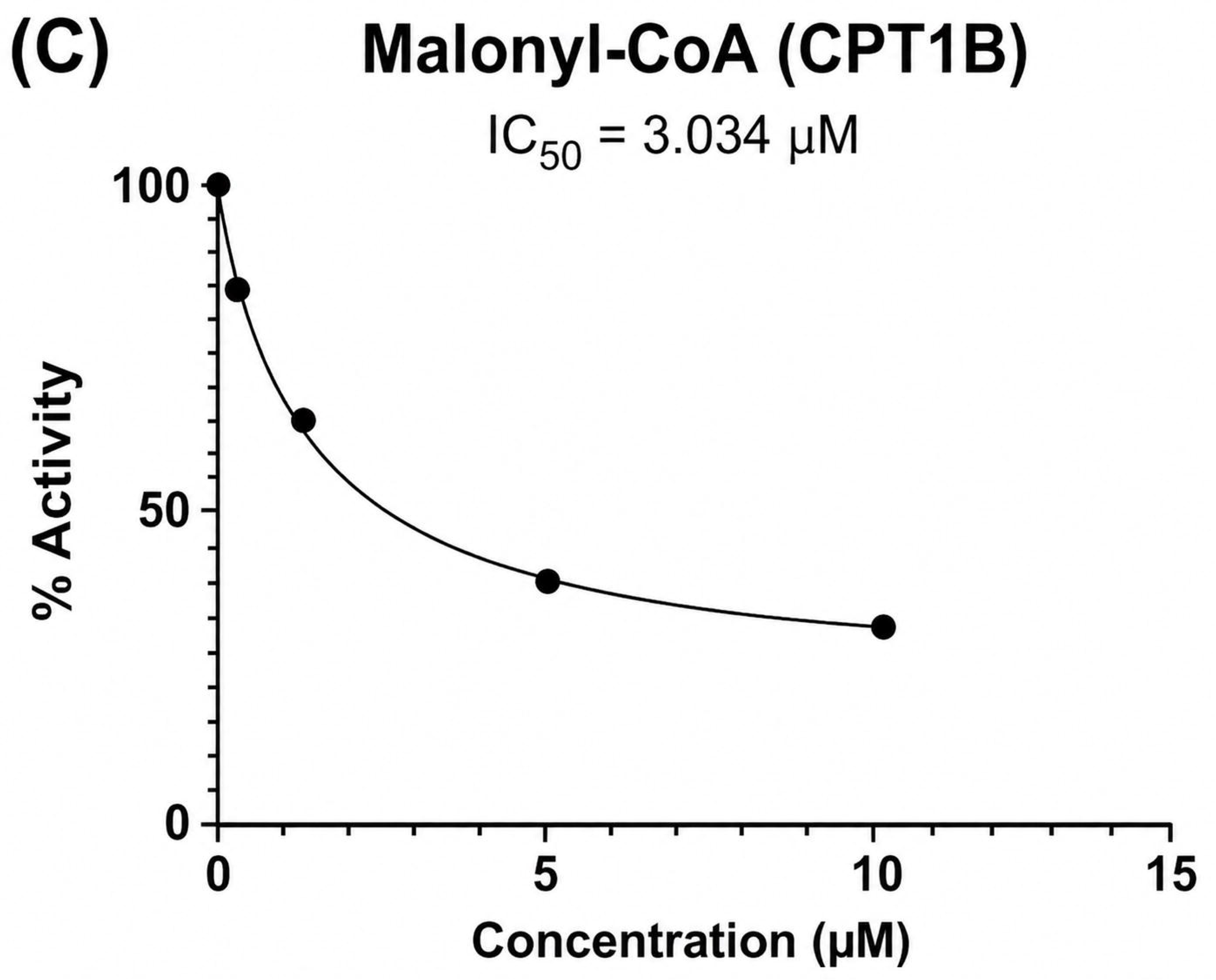
CPT1b inhibitor sensitivity IC_50_ assay with known compounds. Equal amounts of human CPT1b pellet solutions (1.28 μg/μL, 6 μL per triplicate) were tested with a series of inhibitor concentrations (1, 2, 4, 8, and 16 μL) of (R)-(+)-etomoxir (**A**), perhexiline (**B**), and malonyl-CoA (**C**) dissolved in DMSO. DMSO alone (2 μL) served as a negative control. The inhibition curves of (R)-(+)-etomoxir, perhexiline, and malonyl-CoA demonstrate that the CPT1 enzyme activity assay is applicable to both isoforms. (R)-(+)-Etomoxir (**A**) achieved an IC_50_ value of 91.96 nM. This value falls within the acceptable ranges of 10–700 nM (23). Two additional inhibitors, perhexiline with IC_50_ value of 153.2 μM (**B**) and malonyl-CoA with IC_50_ value of 3.034 μM (**C**), were used to validate this platform.

### Identification of CPT1b Inhibitors

To determine further inhibitory or activating effects on CPT1b, small molecules with different indications were prepared and screened. From this screening, we determined that chlorpromazine is a true inhibitor of CPT1b (Figure 4). Notably, CPT1b is more sensitive to inhibitory effects from chlorpromazine than CPT1a, with IC_50_ values of 6.973 µM and 0.1793 mM (19), respectively (Figure 3).

**Figure 4.**
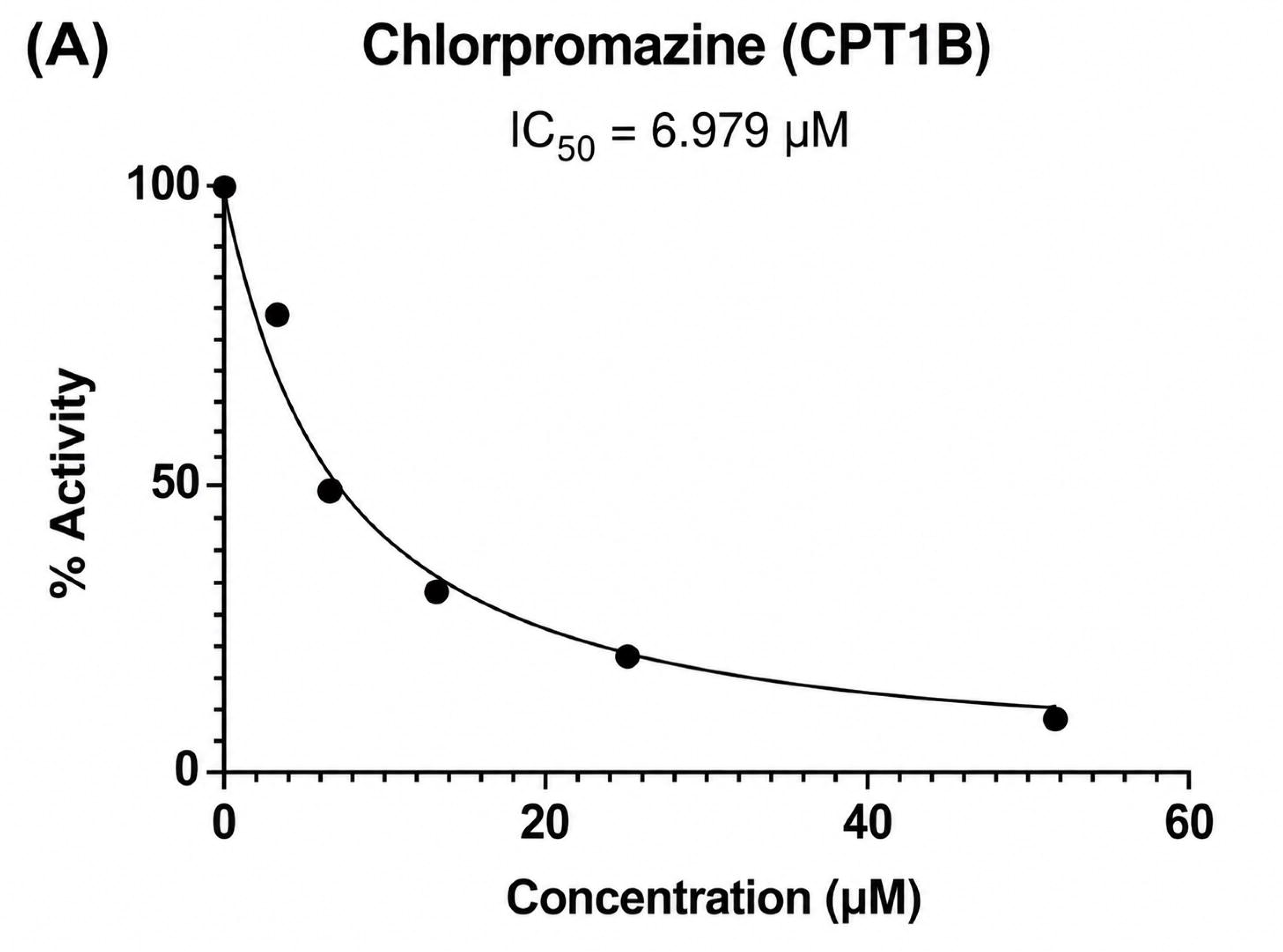
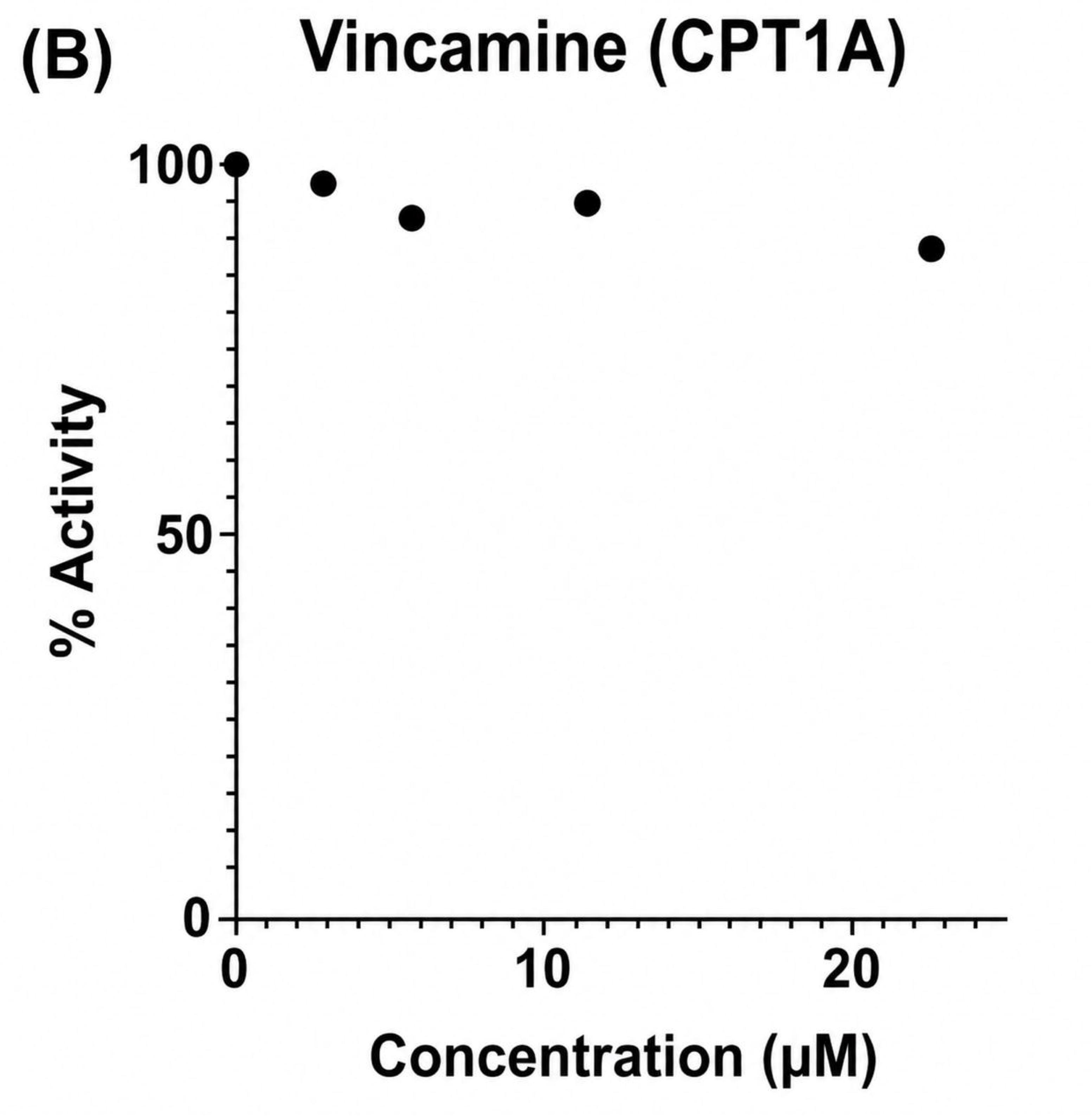
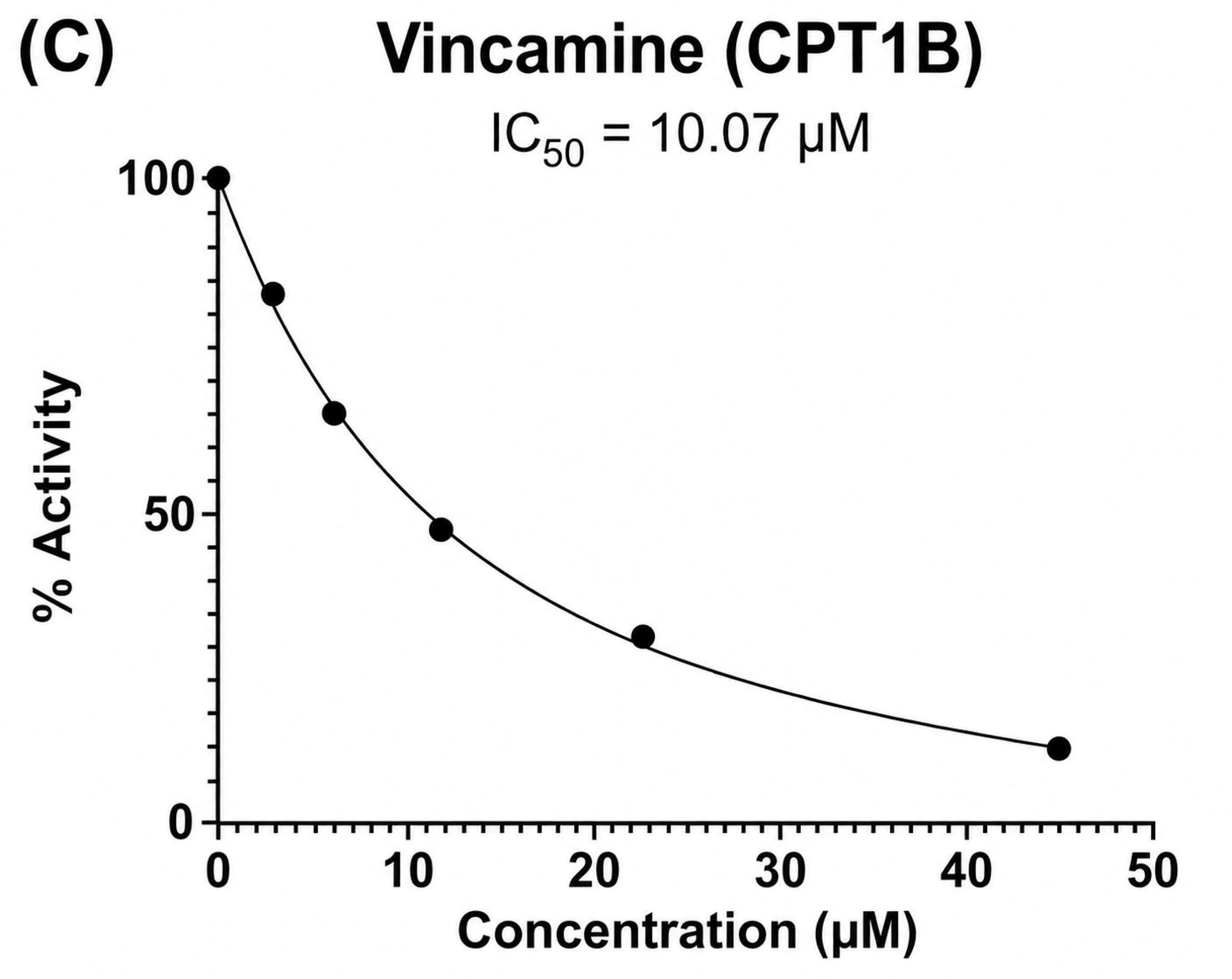
Inhibition of CPT1b by Chlorpromazine and Selective Inhibition of CPT1b by Vincamine. CPT1b inhibitor sensitivity IC_50_ assay with chlorpromazine, achieving an IC_50_ value of 6.979 µM (**A**). Equal amounts of human CPT1b pellet solutions (1.28 μg/μL, 6 μL per triplicate) were tested with a series of inhibitor concentrations (1, 2, 4, 8, and 16 μL) of chlorpromazine. To identify IC₅₀ values of vincamine on CPT1 isoforms dose-response curves were used. Dose–response curves showed (**B**) no inhibition of CPT1a by vincamine and (**C**) inhibition of CPT1b by vincamine having an IC₅₀ value of 10.07 µM. A series of compound concentrations was tested using human CPT1a or CPT1b pellet solutions at 0.67 μg/μL, with 2 μL of protein added per assay.

**Figure 5.**
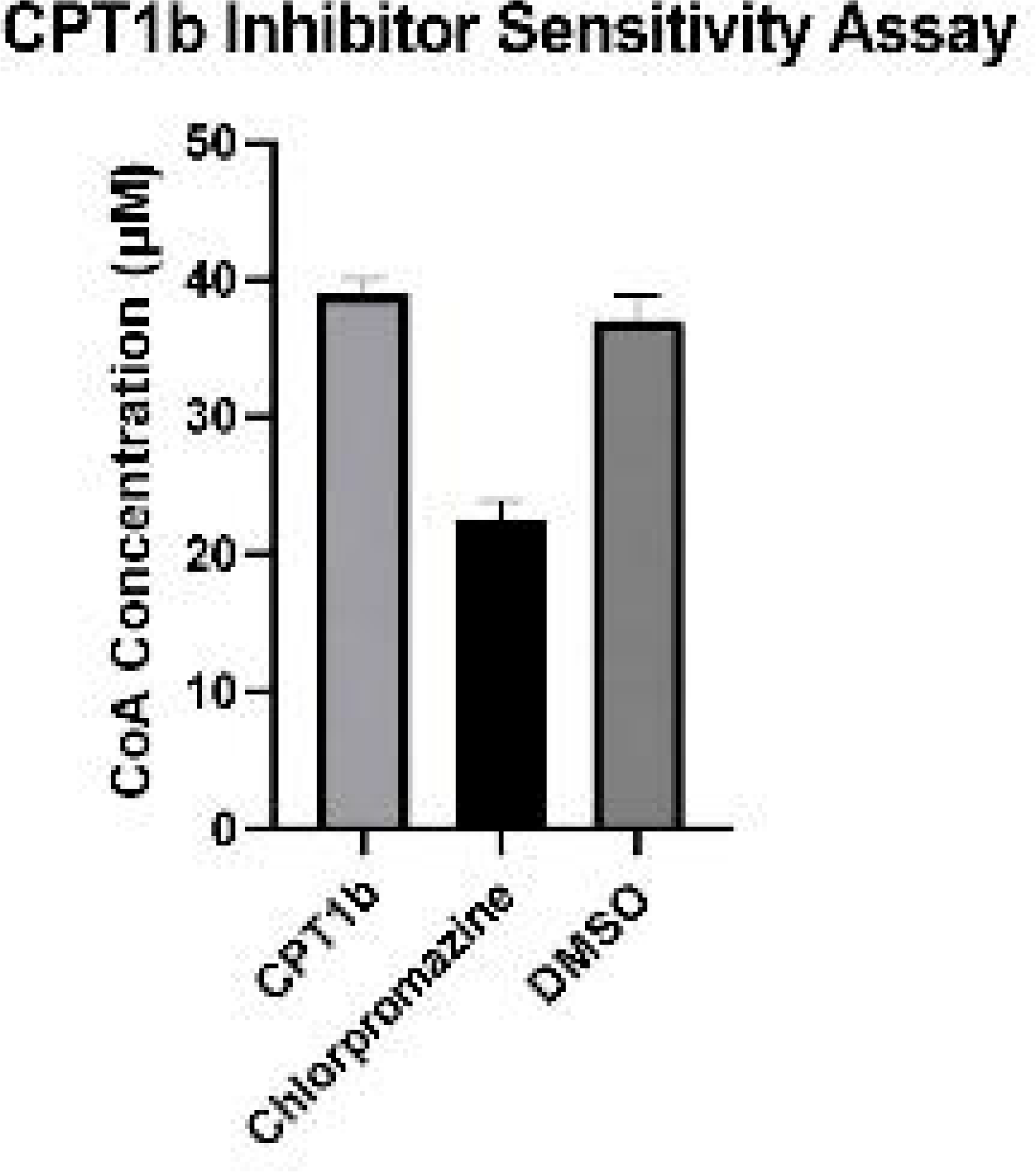
CPT1b Inhibitor Sensitivity Assay. CPT1b activity was compared with a 10mM DMSO control. Enzyme activity was unaffected by DMSO, which compounds are dissolved in. Chlorpromazine showed an inhibitory effect on CPT1b.

To identify isoform-specific inhibitors, CPT1b activity was compared with CPT1a.

Vincamine was discovered to be a CPT1b-selective inhibitor, effectively inhibiting CPT1b while showing no inhibition of CPT1a (Figure 4). Given this selective and potent profile, vincamine emerges as a highly promising candidate drug for targeted CPT1b therapy.

## Discussion

In this study, we utilized a uniform, high-throughput DTNB-based enzyme activity assay platform (19) and successfully adapted it to detect CPT1b enzyme activity and evaluate inhibitor sensitivity. By implementing this established protocol, we enable a side-by-side comparison of inhibitor potency and selectivity between CPT1a and CPT1b.

To assess specificity between CPT1 isoforms, we compared inhibitors using selectivity ratios, defined as IC_50_ (off-target) ÷ IC_50_ (target) (24). A high selectivity ratio (e.g., ≥ 10-fold) implies preferential inhibition for CPT1b. In contrast, ratios approaching 1 suggest broader, non-selective inhibition.

From our screening, we identified one inhibitor that demonstrated selective inhibition for the CPT1b isoform: vincamine. Vincamine was not previously known to be a CPT1 modulator; however, it has been previously reported to show antioxidant activity and cancer cell cytotoxicity by modulating other proteins involved in tumor growth, such as acetylcholinesterase (25). The discovery of vincamine as a selective CPT1b inhibitor therefore represents a mechanism of action distinct from its previously characterized biological targets. Critically, vincamine demonstrated robust inhibition of CPT1b while exhibiting no measurable inhibition of CPT1a, satisfying the ≥10-fold selectivity ratio threshold defined above for isoform-selective inhibitors. Given that CPT1a and CPT1b share only 63% amino acid identity and differ substantially in malonyl-CoA sensitivity and carnitine affinity (3, 5), vincamine’s selectivity is likely attributable to the isoform-specific structural features, though the precise binding interactions remain to be characterized. This selectivity is pharmacologically significant on two levels: first, compounds that spare CPT1a are expected to avoid the hepatotoxicity and oxidative stress associated with broad CPT1 inhibitors such as (R)-(+)-etomoxir and perhexiline (14, 26, 27); second, because CPT1b accounts for approximately 98% of cardiac CPT1 activity (4), selective inhibition of this isoform enables precise modulation of FAO in cardiac and skeletal muscle, the tissues most relevant to metabolic and cardiovascular disease, without disrupting hepatic lipid metabolism (27).

We also demonstrate for the first time that chlorpromazine inhibits CPT1b activity, with an IC_50_ of 6.979 µM (Figure 4). While chlorpromazine has previously been characterized as a broad CPT1 inhibitor and shown to inhibit CPT1a (28), its activity against CPT1b had not been reported. Together with prior literature, this finding completes the known isoform inhibitory profile of chlorpromazine across both CPT1 isoforms and raises questions about whether its known metabolic side effects in clinical use may be partly attributable to CPT1b-mediated suppression of FAO in muscle and cardiac tissue (29).

Selective inhibitors of the various CPT1 isoforms may be of great significance to future therapeutic strategies, minimizing off-target effects associated with broad inhibition of the CPT1 family (30). Previous use of non-specific inhibitors, such as (R)-(+)-etomoxir and perhexiline, has been limited by toxicity, highlighting the need for safer, isoform-targeted therapeutics (14, 26). On the other hand, the CPT1a selective inhibitor ST1326 (Teglicar) improved glycemia and reduced gluconeogenesis in preclinical models (31).

However, the therapeutic potential of CPT1b-selective inhibition in metabolic disease is more complex and context-dependent. Unlike CPT1a inhibition, which directly reduces hepatic glucose production, CPT1b inhibition primarily affects FAO in skeletal and cardiac muscle and may therefore provide limited benefit in improving systemic glycemic control (32, 33). While acute reductions in FAO can briefly enhance glucose utilization, chronic inhibition has been associated with the accumulation of lipotoxic intermediates and may exacerbate insulin resistance in metabolic tissues (34, 35).

Moreover, given that CPT1b accounts for 98% of CPT1 activity in cardiac tissue (4), sustained inhibition of this isoform may disrupt myocardial energy homeostasis. The adult heart relies primarily on FAO for ATP production, and impairment of this pathway has been linked to metabolic remodeling, lipid accumulation, and pathological cardiac hypertrophy (32, 37). Thus, although CPT1b-selective inhibitors may avoid the hepatotoxicity associated with CPT1a inhibition (14, 26, 27), they may introduce a distinct risk of cardiovascular dysfunction by localizing metabolic stress to cardiac tissue (38, 39).

In contrast, CPT1b inhibition may be more favorably applied in oncological contexts, where suppression of FAO can limit energy production in tumors that rely on oxidative metabolism for growth (40, 41). CPT1b-selective inhibitors could therefore possibly serve as targeted metabolic modulators in cancer while requiring careful evaluation of long-term safety in metabolic and cardiovascular disease settings (42).

Ultimately, the development of isoform-selective CPT1b inhibitors represents a promising opportunity for achieving efficacy with improved safety profiles.

Future work could include improving assay sensitivity and enzyme validation to address current limitations. Fluorescence-based assays and additional protein characterization using SDS-PAGE and Western blot would enhance confidence in inhibitor screening results (43, 44).

Factors such as substrate concentration and enzyme preparation can influence absolute IC_50_ values (45). High-throughput biochemical assays may employ non-physiological substrate concentrations or detection systems that differ from native mitochondrial physiological conditions, subtly affecting apparent inhibitor potency (46). Furthermore, the use of mammalian cell-derived protein rather than purified recombinant enzymes introduces inherent biological volatility (47). Residual host cell proteins and cellular components present in mammalian expression systems can lead to non-specific binding or interfere with enzymatic measurements, potentially increasing the observed IC_50_ relative to purified recombinant systems (48). Additionally, recombinant expression of mitochondrial proteins involved in FAO, including CPT1b, enables efficient high-throughput screening but may not fully replicate native mitochondrial folding or post-translational modifications, which could subtly influence inhibitor binding or enzymatic activity (49, 50).

Lastly, inhibitor potency in vitro does not always predict biological activity (51). In cellular or in vivo contexts, factors such as membrane permeability, metabolic stability, protein binding, and subcellular distribution can substantially affect the extent to which an inhibitor engages its target (52).

In summary, our DTNB-based high-throughput assay enables the efficient identification of small-molecule inhibitors of CPT1b, providing a reliable platform to evaluate compound efficacy and enzyme activity. Subsequent trials across CPT1 isoforms revealed inhibitors with distinct isoform-selective activity, an important feature given the tissue-specific expression of these isoforms. Chlorpromazine and vincamine are both CPT1b inhibitors, and vincamine demonstrates strong inhibition of CPT1b while showing no detectable activity against CPT1a. The isoform selectivity of vincamine can minimize off-target metabolic effects and systemic toxicity, providing a basis for isoform-specific side effect mapping and guiding safer, more targeted therapeutic interventions. The precise targeting of CPT1b by vincamine can help modulate lipid oxidation in specific tissues for metabolic diseases, making it a candidate drug for cancer (17), metabolic disorders (35), and cardiovascular conditions (37). Therefore, our platform demonstrates the potential of isoform-selective inhibition as a strategy for the development of precise, safe, and effective treatments in metabolic and oncological disorders.

## Experimental Procedures

### Cell Culture and CPT1 Transfection

Mammalian Expi293F cells (Gibco, WA, USA; Cat. #A14527) were stably transfected with a human CPT1b expression plasmid using the ExpiFectamine 293 Kit (Gibco, WA, USA; Cat. #A14527), and a parallel Expi293F line was generated by stable transfection with a human CPT1a expression plasmid to enable comparative analysis of isoform-specific differences. All cells were maintained in Expi293 Expression Medium (Gibco, WA, USA, Cat. #12338018) and cultured in 250 mL bottles in a humidified incubator at 37°C with 5% CO_2_.

Expression plasmids encoding human CPT1a and CPT1b were constructed and processed using parallel workflows. For CPT1a, two constructs—CPT1a (NM_001876.3) ORF containing the full-length human coding sequence, and sp_CPT1a (NM_001876.3) ORF containing an N-terminal signal peptide for extracellular release—were cloned into the pcDNA3.1+/C-(K)-DYK vector backbone and obtained from GenScript USA Inc. (Piscataway, NJ, USA). The CPT1b expression plasmid was generated using the CPT1b ORF reference sequence (see Supporting Information), containing the full-length human coding sequence, with an optional CPT1b ORF including an N-terminal signal peptide for extracellular release if applicable. All plasmids were transformed into TOP10 bacteria for amplification, followed by plasmid extraction, purification, and Sanger sequencing to verify correct insertion of the respective CPT1a and CPT1b genes. Verified plasmids were then transfected into mammalian Expi293F cell lines. After expansion of Expi293F cultures, both the supernatant and cell pellets were lysed using RIPA buffer [25 mM Tris·HCl pH 7.6, 150 mM NaCl, 1% NP-40, 1% sodium deoxycholate, 0.1% SDS] (ThermoFisher, Bothell, WA, USA; Cat. #88901). Proteins were harvested through differential, two-step centrifugation modified from Yang et al. (53), with the remaining mitochondrial pellet resuspended in the extraction buffer (54). Protein concentrations were quantified using a Bradford protein assay kit.

### Mitochondrial Marking for Normalization

To normalize mitochondrial protein loading and confirm mitochondrial enrichment, a TOM20 recombinant polyclonal antibody (Cat # 11802-1-AP, 150 uL, Proteintech) was used as a mitochondrial outer membrane marker to quantify the relative expression of CPT1a and CPT1b. Because TOM20, CPT1a, and CPT1b are localized in the mitochondrial outer membrane, TOM20 serves as an appropriate within-compartment reference protein and normalization control for mitochondrial protein assays. Normalizing CPT1a and CPT1b expression to TOM20 ensures that equal amounts of mitochondrial protein are compared across samples, accounting for variability in mitochondrial isolation efficiency and transfection between controls, with isolated mitochondrial samples generally containing more protein than nontransgene samples. This ensures that observed changes in CPT1a and CPT1b expression reflect true biological differences rather than technical variability introduced during transfection, mitochondrial extraction, or immunoassay preparation. Additionally, normalization to total protein concentration via Bradford assay ensures that CPT1a and CPT1b activity measurements are directly comparable across mitochondrial extracts. Goat-anti-rabbit HRP secondary antibody (ThermoFisher, Bothell, WA, USA; Cat. #31460) was used in tandem with the TOM20 recombinant polyclonal antibody.

### Immunoassay Quantification and Normalization of CPT1a and CPT1b

Enzyme-linked immunosorbent assay (ELISA) was used to detect the expression level of human CPT1a and CPT1b in the supernatant and pellet and to confirm successful transgenic expression. The CPT1b-transfected and CPT1a-transfected cell pellets and supernatants, containing 1 µg of total protein, were coated in freshly prepared NaHCO_3_-Na_2_CO_3_ buffer (pH 6.4) and incubated at 4°C for 18 hours. Following three washes with PBST (PBS + 0.1% Tween-20; 100 μL/wash), 200 µL of blocking buffer (2.5% nonfat milk in PBST) was added to each well for 1 hour at 18°C. Mouse anti-human CPT1b monoclonal antibody (ThermoFisher, Bothell, WA, USA; Cat. #MA5-27329) or mouse anti-human CPT1a monoclonal antibody (Abcam, Fremont, CA, USA; Cat. #8F6AE9), depending on the construct of interest, was added in 1:1000 dilution and shaken for 2 hours at 18°C. The goat-anti-mouse horseradish peroxidase (HRP) secondary antibody (Rockland Immunochemicals, Pottstown, PA, USA; Cat. #610-1303) was added with the blocking buffer after a second wash. 100 µL of the diluted secondary antibody was then added to each well after washing three times with PBST for 1 hour at 18°C while shaking. Then, 50 µL 3,3’,5,5’-Tetramethylbenzidine substrate (0.4 g/L, Tri\bioscience, Sunnyvale, CA, USA; Cat. #TBS5021) was added to each well after three washes of PBST at 18°C. A total of 100 µL of 1 M H_2_SO_4_ stop solution was then added to each well. Absorbance was measured at 450 nm after 10 seconds of shaking, with Expi293F cell pellet protein as an endogenous control.

### CPT Enzyme Activity Assay and Inhibitor Sensitivity Assay

Compounds were prepared at 10 mM in dimethyl sulfoxide (DMSO; Stellar Chemical), which had no detectable effect on assay performance. Perhexiline, malonyl-CoA, and (R)-(+)-etomoxir were included as positive controls. Varying concentrations of (R)-(+)-etomoxir, adjusted with proportional DMSO volumes, were tested to determine optimal assay sensitivity. The enzyme volume was optimized by assessing multiple concentrations for the strongest signal-to-background response. To account for a high background signal and normalize activity, the absorbance value obtained from heat-denatured enzyme controls was subtracted from all experimental readings. Prior to compound screening, a standard curve was generated to confirm linearity between absorbance at 412 nm and enzymatic reaction rate under assay conditions.

This method was a slight modification to previously established DTNB-based assays (26, 27). Compounds were added to the designated wells and mixed, followed by a pre-prepared, thawed aliquot of the CPT1b-containing mitochondrial extract to each well. Reactions were incubated at 30°C for 10 minutes to maximize enzyme activity, after which 10 μL of 1 mg/mL DTNB (diluted in 100 mM tris buffer) was added. Plates were gently shaken using a plate shaker at 400 revolutions per minute (rpm) for 10 minutes to ensure thorough mixing and then read for absorbance at 412 nm - 600 nm.

### Identification and Characterization of Selective Inhibitors

To assess the suitability of this platform for large-scale, high-throughput screening, a library of small-molecule compounds, drawn from a diverse collection of natural products, antivirals, immunomodulators, and other bioactive molecules, was applied, with (R)-(+)-etomoxir used to validate the assay. Compounds were diluted to 10 mM in dimethyl sulfoxide (Stellar Chemical, Rahway, NJ, USA, 99.99%) and tested at a final concentration of 0.556 mM. All assays were performed in 96-well plates in triplicate, with control wells included to quantify baseline enzyme activity.

Compounds that produced significant inhibition relative to controls were further evaluated by IC_50_ determination using serial doubling dilutions performed in triplicate at a fixed substrate concentration under standard DTNB assay conditions. Dose–response curves were generated for both CPT1a and CPT1b to assess isoform selectivity, and enzyme concentrations were normalized using a Bradford assay to optimize signal-to-background ratios. Selective inhibitors were defined as compounds demonstrating differential inhibition between CPT1a and CPT1b.

### Data Analysis

All statistical analyses (means, error values, and t-tests) were carried out using GraphPad Prism 10. Results are shown as mean ± SEM (n=3), and differences were considered significant when p < 0.05. Dose-response curves were analyzed using a three parameter nonlinear regression model ([inhibitor] vs response), with the top constrained to maximal observed response.

### Materials

Assays were performed with a mastermix solution composed of 1.5 mM Ethylenediaminetetraacetic acid (Acros, Geel, Belgium, Cat. #AC118432500), 40 mM 4-(2-Hydroxyethyl)-1-piperazine ethanesulfonic acid (ThermoScientific, Waltham, MA, USA, Cat. #15630080), 35 μM Palmitoyl-CoA (Sigma Aldrich, St. Louis, MO, USA, Cat. #P9716), 1.25 mM β-hydroxy-γ-N-trimethyl aminobutyric acid (AK Scientific, Union City, NJ, USA, Cat. #J94077), and 100 mM of Tris buffer to equalize well volumes to 90 μL (19).

## Data Availability

The raw data supporting the conclusions of this article will be made available by the authors upon request, addressed to the contact authors.

## Supporting Information

This article contains supporting information.

## Conflict of interest

The authors declare that they have no conflicts of interest with the contents of this article.

## Acknowledgments

The authors acknowledge the support of the Aspiring Scholars Directed Research Program (ASDRP).

## Author contributions

A.W., W.L., J.X., H.G., M.L., A.N., S.M., H.T., C.W., M.B., and M.W.: conceptualization, data curation, formal analysis, investigation, methodology, validation, writing–original draft, and writing–review & editing; Z.C.: conceptualization, supervision, project administration, resources, funding acquisition, and writing–review & editing.

## Funding and additional information

This study was supported and funded by the Olive Children Foundation.

## Abbreviations

CPT1: carnitine palmitoyltransferase 1
CPT1a: carnitine palmitoyltransferase 1a
CPT1b: carnitine palmitoyltransferase 1b
CPT1c: carnitine palmitoyltransferase 1c
CoA: coenzyme A
DTNB: 5,5′-Dithiobis(2-nitrobenzoic acid)
ELISA: enzyme-linked immunosorbent assay
HRP: horseradish peroxidase
TCA cycle: tricarboxylic acid cycle
TNB: 1,3,5-Trinitrobenzene.

## Notes

### Competing Interest Statement

The authors have declared no competing interest.

### Summary of Updates

The ORCID id between authors 2 and 4 were mixed up. The numbers were updated accordingly.

