## Supporting Information for "Comparative profiling of carnitine palmitoyltransferase 1 isoforms reveals vincamine as a selective carnitine palmitoyltransferase 1b inhibitor"

A. CPT1a Non-fusion Plasmid S-1
   a. Non-fusion Plasmid Sequence S-1
   b. Figure S1: Non-fusion Plasmid Map S-2
B. CPT1a Fusion Plasmid S-3
   a. Fusion Plasmid Sequence S-3
   b. Figure S2: Fusion Plasmid Map S-6
C. CPT1b Non-fusion Plasmid S-7
   a. Non-fusion Plasmid Sequence S-7
   b. Figure S3: Non-fusion Plasmid Map S-10
D. CPT1b Fusion Plasmid S-11
   a. Fusion Plasmid Sequence S-11
   b. Figure S4: Fusion Plasmid Map S-12
E. Figure S5: Standard Curve for DTNB S-13
F. Figure S6: Bradford S-13

### A. CPT1A Non-fusion Plasmid

#### a. Non-fusion Plasmid Sequence

pcDNA3.1

> CPT1A(NM_001876.3) ORF Clone

ATGGCAGAAGCTCACCAAGCTGTGGCCTTTCAGTTCACGGTCACTCCGGACGGGATTGACCT
GCGGCTGAGCCATGAAGCTCTTAGACAAATCTATCTCTCTGGACTTCATTCCTGGAAAAAGAA
GTTCATCAGATTCAAGAACGGCATCATCACTGGCGTGTACCCGGCAAGCCCCTCCAGTTGGCT
TATCGTGGTGGTGGGCGTGATGACAACGATGTACGCCAAGATCGACCCCTCGTTAGGAATAAT
TGCAAAAATCAATCGGACTCTGGAAACGGCCAACTGCATGTCCAGCCAGACGAAGAACGTGG
TCAGCGGCGTGCTGTTTGGCACCGGCCTGTGGGTGGCCCTCATCGTCACCATGCGCTACTCCC
TGAAAGTGCTGCTCTCCTACCACGGGTGGATGTTCACTGAGCACGGCAAGATGAGTCGTGCC
ACCAAGATCTGGATGGGTATGGTCAAGATCTTTTCAGGCCGAAAACCCATGTTGTACAGCTTC
CAGACATCGCTGCCTCGCCTGCCGGTCCCGGCTGTCAAAGACACTGTGAACAGGTATCTACA
GTCGGTGAGGCCTCTTATGAAGGAAGAAGACTTCAAACGGATGACAGCACTTGCTCAAGATT
TTGCTGTCGGTCTTGGACCAAGATTACAGTGGTATTTGAAGTTAAAATCCTGGTGGGCTACAA
ATTACGTGAGCGACTGGTGGGAGGAGTACATCTACCTCCGAGGACGAGGGCCGCTCATGGTG
AACAGCAACTATTATGCCATGGATCTGCTGTATATCCTTCCAACTCACATTCAGGCAGCAAGAG
CCGGCAACGCCATCCATGCCATCCTGCTTTACAGGCGCAAACTGGACCGGGAGGAAATCAAA
CCAATTCGTCTTTTGGGATCCACGATTCCACTCTGCTCCGCTCAGTGGGAGCGGATGTTTAATA
CTTCCCGGATCCCAGGAGAGGAGACAGACACCATCCAGCACATGAGAGACAGCAAGCACATC
GTCGTGTACCATCGAGGACGCTACTTCAAGGTCTGGCTCTACCATGATGGGCGGCTGCTGAAG
CCCCGGGAGATGGAGCAGCAGATGCAGAGGATCCTGGACAATACCTCGGAGCCTCAGCCCGG
GGAGGCCAGGCTGGCAGCCCTCACCGCAGGAGACAGAGTTCCCTGGGCCAGGTGTCGTCAG
GCCTATTTTGGACGTGGGAAAAATAAGCAGTCTCTTGATGCTGTGGAGAAAGCAGCGTTCTTC
GTGACGTTAGATGAAACTGAAGAAGGATACAGAAGTGAAGACCCGGATACGTCAATGGACAG
CTACGCCAAATCTCTACTACACGGCCGATGTTACGACAGGTGGTTTGACAAGTCGTTCACGTT
TGTTGTCTTCAAAAACGGGAAGATGGGCCTCAACGCTGAACACTCCTGGGCAGATGCGCCGA
TCGTGGCCCACCTTTGGGAGTACGTCATGTCCATTGACAGCCTCCAGCTGGGCTATGCGGAGG
ATGGGCACTGCAAAGGCGACATCAATCCGAACATTCCGTACCCCACCAGGCTGCAGTGGGAC
ATCCCGGGGGAATGTCAAGAGGTTATAGAGACCTCCCTGAACACCGCAAATCTTCTGGCAAA
CGACGTGGATTTCCATTCCTTCCCATTCGTAGCCTTTGGTAAAGGAATCATCAAGAAATGTCGC
ACGAGCCCAGACGCCTTTGTGCAGCTGGCCCTCCAGCTGGCGCACTACAAGGACATGGGCAA
GTTTTGCCTCACATACGAGGCCTCCATGACCCGGCTCTTCCGAGAGGGGAGGACGGAGACCG
TGCGCTCCTGCACCACTGAGTCATGCGACTTCGTGCGGGCCATGGTGGACCCGGCCCAGACG
GTGGAACAGAGGCTGAAGTTGTTCAAGTTGGCGTCTGAGAAGCATCAGCATATGTATCGCCTC
GCCATGACCGGCTCTGGGATCGATCGTCACCTCTTCTGCCTTTACGTGGTGTCTAAATATCTCG
CTGTGGAGTCCCCTTTCCTTAAGGAAGTTTTATCTGAGCCTTGGAGATTATCAACAAGCCAGA
CCCCTCAGCAGCAAGTGGAGCTGTTTGACTTGGAGAATAACCCAGAGTACGTGTCCAGCGGA
GGGGGCTTTGGACCGGTTGCTGATGACGGCTATGGTGTGTCGTACATCCTTGTGGGAGAGAAC
CTCATCAATTTCCACATTTCTTCCAAGTTCTCTTGCCCTGAGACGGATTCTCATCGCTTTGGAA
GGCACCTGAAAGAAGCAATGACTGACATCATCACTTTGTTTGGTCTCAGTTCTAATTCCAAAA
AG

b. **Figure S1: Non-fusion Plasmid Map**
**After the insertion following common subclone protocols, we inserted our CPT1a wild-type gene into our pcDNA 3.1 vector into the cloning MCS site. After we sequenced, there is no new mutation, we validated the start point and the stop codon works well. There is no reading frameshift mutation. We then used this vector for our CPT1a and CPT1b protein harvest.**

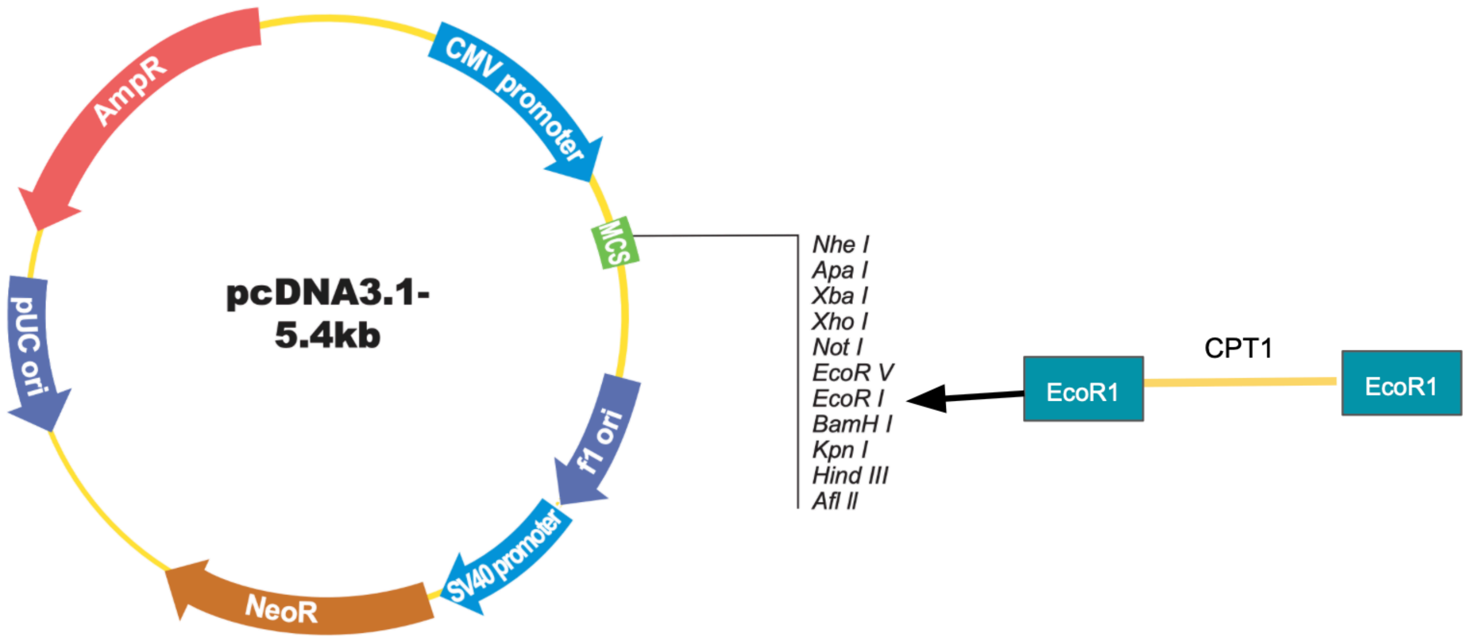

### B. CPT1A Fusion Plasmid

#### a. Fusion Plasmid Sequence

>CPT1A(NM_001876.3) ORF Clone
GACGGATCGGGAGATCTCCCGATCCCCTATGGTGCACTCTCAGTACAATCTGCTCTGATGCCG
CATAGTTAAGCCAGTATCTGCTCCCTGCTTGTGTGTTGGAGGTCGCTGAGTAGTGCGCGAGCA
AAATTTAAGCTACAACAAGGCAAGGCTTGACCGACAATTGCATGAAGAATCTGCTTAGGGTTA
GGCGTTTTGCGCTGCTTCGCGATGTACGGGCCAGATATACGCGTTGACATTGATTATTGACTAG
TTATTAATAGTAATCAATTACGGGGTCATTAGTTCATAGCCCATATATGGAGTTCCGCGTTACATA
ACTTACGGTAAATGGCCCGCCTGGCTGACCGCCCAACGACCCCCGCCCATTGACGTCAATAAT
GACGTATGTTCCCATAGTAACGCCAATAGGGACTTTCCATTGACGTCAATGGGTGGAGTATTTA
CGGTAAACTGCCCACTTGGCAGTACATCAAGTGTATCATATGCCAAGTACGCCCCCTATTGACG
TCAATGACGGTAAATGGCCCGCCTGGCATTATGCCCAGTACATGACCTTATGGGACTTTCCTAC
TTGGCAGTACATCTACGTATTAGTCATCGCTATTACCATGGTGATGCGGTTTTGGCAGTACATCA
ATGGGCGTGGATAGCGGTTTGACTCACGGGGATTTCCAAGTCTCCACCCCATTGACGTCAATG
GGAGTTTGTTTTGGCACCAAAATCAACGGGACTTTCCAAAATGTCGTAACAACTCCGCCCCAT
TGACGCAAATGGGCGGTAGGCGTGTACGGTGGGAGGTCTATATAAGCAGAGCTCTCTGGCTAA
CTAGAGAACCCACTGCTTACTGGCTTATCGAAATTAATACGACTCACTATAGGGAGACCCAAG
CTGGCTAGCGTTTAAACTTAAGCTTGGTACCGAGCTCGGATCCGCCACCATGGCTGGAGCTG
CATCATCCTGTTCCTGGTGGCCACAGCCACCGGCGTGCACTCTGCAGAAGCTCACCAAGCTGT
GGCCTTTCAGTTCACGGTCACTCCGGACGGGATTGACCTGCGGCTGAGCCATGAAGCTCTTAG
ACAAATCTATCTCTCTGGACTTCATTCCTGGAAAAAGAAGTTCATCAGATTCAAGAACGGCAT
CATCACTGGCGTGTACCCGGCAAGCCCCTCCAGTTGGCTTATCGTGGTGGTGGGCGTGATGAC
AACGATGTACGCCAAGATCGACCCCTCGTTAGGAATAATTGCAAAAATCAATCGGACTCTGGA
AACGGCCAACTGCATGTCCAGCCAGACGAAGAACGTGGTCAGCGGCGTGCTGTTTGGCACCG
GCCTGTGGGTGGCCCTCATCGTCACCATGCGCTACTCCCTGAAAGTGCTGCTCTCCTACCACG
GGTGGATGTTCACTGAGCACGGCAAGATGAGTCGTGCCACCAAGATCTGGATGGGTATGGTC
AAGATCTTTTCAGGCCGAAAACCCATGTTGTACAGCTTCCAGACATCGCTGCCTCGCCTGCCG
GTCCCGGCTGTCAAAGACACTGTGAACAGGTATCTACAGTCGGTGAGGCCTCTTATGAAGGA
AGAAGACTTCAAACGGATGACAGCACTTGCTCAAGATTTTGCTGTCGGTCTTGGACCAAGATT
ACAGTGGTATTTGAAGTTAAAATCCTGGTGGGCTACAAATTACGTGAGCGACTGGTGGGAGG
AGTACATCTACCTCCGAGGACGAGGGCCGCTCATGGTGAACAGCAACTATTATGCCATGGATC
TGCTGTATATCCTTCCAACTCACATTCAGGCAGCAAGAGCCGGCAACGCCATCCATGCCATCC
TGCTTTACAGGCGCAAACTGGACCGGGAGGAAATCAAACCAATTCGTCTTTTGGGATCCACG
ATTCCACTCTGCTCCGCTCAGTGGGAGCGGATGTTTAATACTTCCCGGATCCCAGGAGAGGAG
ACAGACACCATCCAGCACATGAGAGACAGCAAGCACATCGTCGTGTACCATCGAGGACGCTA
CTTCAAGGTCTGGCTCTACCATGATGGGCGGCTGCTGAAGCCCCGGGAGATGGAGCAGCAGA
TGCAGAGGATCCTGGACAATACCTCGGAGCCTCAGCCCGGGGAGGCCAGGCTGGCAGCCCTC
ACCGCAGGAGACAGAGTTCCCTGGGCCAGGTGTCGTCAGGCCTATTTTGGACGTGGGAAAAA
TAAGCAGTCTCTTGATGCTGTGGAGAAAGCAGCGTTCTTCGTGACGTTAGATGAAACTGAAG
AAGGATACAGAAGTGAAGACCCGGATACGTCAATGGACAGCTACGCCAAATCTCTACTACAC
GGCCGATGTTACGACAGGTGGTTTGACAAGTCGTTCACGTTTGTTGTCTTCAAAAACGGAA
GATGGGCCTCAACGCTGAACACTCCTGGGCAGATGCGCCGATCGTGGCCCACCTTTGGGAGT

ACGTCATGTCCATTGACAGCCTCCAGCTGGGCTATGCGGAGGATGGGCACTGCAAAGGCGAC
ATCAATCCGAACATTCCGTACCCCACCAGGCTGCAGTGGGACATCCCGGGGGAATGTCAAGA
GGTTATAGAGACCTCCCTGAACACCGCAAATCTTCTGGCAAACGACGTGGATTTCCATTCCTT
CCCATTCGTAGCCTTTGGTAAAGGAATCATCAAGAAATGTCGCACGAGCCCAGACGCCTTTGT
GCAGCTGGCCCTCCAGCTGGCGCACTACAAGGACATGGGCAAGTTTTGCCTCACATACGAGG
CCTCCATGACCCGGCTCTTCCGAGAGGGGAGGACGGAGACCGTGCGCTCCTGCACCACTGAG
TCATGCGACTTCGTGCGGGCCATGGTGGACCCGGCCCAGACGGTGGAACAGAGGCTGAAGTT
GTTCAAGTTGGCGTCTGAGAAGCATCAGCATATGTATCGCCTCGCCATGACCGGCTCTGGGAT
CGATCGTCACCTCTTCTGCCTTTACGTGGTGTCTAAATATCTCGCTGTGGAGTCCCCTTTCCTTA
AGGAAGTTTTATCTGAGCCTTGGAGATTATCAACAAGCCAGACCCCTCAGCAGCAAGTGGAG
CTGTTTGACTTGGAGAATAACCCAGAGTACGTGTCCAGCGGAGGGGGCTTTGGACCGGTTGC
TGATGACGGCTATGGTGTGTCGTACATCCTTGTGGGAGAGAACCTCATCAATTTCCACATTTCT
TCCAAGTTCTCTTGCCCTGAGACGGATTCTCATCGCTTTGGAAGGCACCTGAAAGAAGCAATG
ACTGACATCATCACTTTGTTTGGTCTCAGTTCTAATTCCAAAAAGGATTACAAGGATGACGAC
GATAAGTGATAAACCCGCTGATCAGCCTCGACTGTGCCTTCTAGTTGCCAGCCATCTGTTGTTT
GCCCCTCCCCCGTGCCTTCCTTGACCCTGGAAGGTGCCACTCCCACTGTCCTTTCCTAATAAA
ATGAGGAAATTGCATCGCATTGTCTGAGTAGGTGTCATTCTATTCTGGGGGGTGGGGTGGGGC
AGGACAGCAAGGGGGAGGATTGGGAAGACAATAGCAGGCATGCTGGGGATGCGGTGGGCTC
TATGGCTTCTGAGGCGGAAAGAACCAGCTGGGGCTCTAGGGGGTATCCCCACGCGCCCTGTA
GCGGCGCATTAAGCGCGGCGGGTGTGGTGGTTACGCGCAGCGTGACCGCTACACTTGCCAGC
GCCCTAGCGCCCGCTCCTTTCGCTTTCTTCCCTTCCTTTCTCGCCACGTTCGCCGGCTTTCCCC
GTCAAGCTCTAAATCGGGGGCTCCCTTTAGGGTTCCGATTTAGTGCTTTACGGCACCTCGACC
CCAAAAAACTTGATTAGGGTGATGGTTCACGTAGTGGGCCATCGCCCTGATAGACGGTTTTTC
GCCCTTTGACGTTGGAGTCCACGTTCTTTAATAGTGGACTCTTGTTCCAAACTGGAACAACAC
TCAACCCTATCTCGGTCTATTCTTTTGATTTATAAGGGATTTTGCCGATTTCGGCCTATTGGTTA
AAAAATGAGCTGATTTAACAAAAATTTAACGCGAATTAATTCTGTGGAATGTGTGTCAGTTAG
GGTGTGGAAAGTCCCCAGGCTCCCCAGCAGGCAGAAGTATGCAAAGCATGCATCTCAATTAG
TCAGCAACCAGGTGTGGAAAGTCCCCAGGCTCCCCAGCAGGCAGAAGTATGCAAAGCATGC
ATCTCAATTAGTCAGCAACCATAGTCCCGCCCCTAACTCCGCCCATCCCGCCCCTAACTCCGCC
CAGTTCCGCCCATTCTCCGCCCCATGGCTGACTAATTTTTTTTATTTATGCAGAGGCCGAGGCC
GCCTCTGCCTCTGAGCTATTCCAGAAGTAGTGAGGAGGCTTTTTTGGAGGCCTAGGCTTTTGC
AAAAAGCTCCCGGGAGCTTGTATATCCATTTTCGGATCTGATCAAGAGACAGGATGAGGATCG
TTTCGCATGATTGAACAAGATGGATTGCACGCAGGTTCTCCGGCCGCTTGGGTGGAGAGGCTA
TTCGGCTATGACTGGGCACAACAGACAATCGGCTGCTCTGATGCCGCCGTGTTCCGGCTGTCA
GCGCAGGGGCGCCCGGTTCTTTTTGTCAAGACCGACCTGTCCGGTGCCCTGAATGAACTGCA
GGACGAGGCAGCGCGGCTATCGTGGCTGGCCACGACGGGCGTTCCTTGCGCAGCTGTGCTCG
ACGTTGTCACTGAAGCGGGAAGGGACTGGCTGCTATTGGGCGAAGTGCCGGGGCAGGATCTC
CTGTCATCTCACCTTGCTCCTGCCGAGAAAGTATCCATCATGGCTGATGCAATGCGGCGGCTG
CATACGCTTGATCCGGCTACCTGCCCATTCGACCACCAAGCGAAACATCGCATCGAGCGAGCA
CGTACTCGGATGGAAGCCGGTCTTGTCGATCAGGATGATCTGGACGAAGAGCATCAGGGGCT
CGCGCCAGCCGAACTGTTCGCCAGGCTCAAGGCGCGCATGCCCGACGGCGAGGATCTCGTCG
TGACCCATGGCGATGCCTGCTTGCCGAATATCATGGTGGAAAATGGCCGCTTTTCTGGATTCAT
CGACTGTGGCCGGCTGGGTGTGGCGGACCGCTATCAGGACATAGCGTTGGCTACCCGTGATAT
TGCTGAAGAGCTTGGCGGCGAATGGGCTGACCGCTTCCTCGTGCTTTACGGTATCGCCGCTCC

CGATTCGCAGCGCATCGCCTTCTATCGCCTTCTTGACGAGTTCTTCTGAGCGGGACTCTGGGG
TTCGAAATGACCGACCAAGCGACGCCCAACCTGCCATCACGAGATTTCGATTCCACCGCCGCC
TTCTATGAAAGGTTGGGCTTCGGAATCGTTTTCCGGGACGCCGGCTGGATGATCCTCCAGCGC
GGGGATCTCATGCTGGAGTTCTTCGCCCACCCCAACTTGTTTATTGCAGCTTATAATGGTTACA
AATAAAGCAATAGCATCACAAATTTCACAAATAAAGCATTTTTTTCACTGCATTCTAGTTGTGG
TTTGTCCAAACTCATCAATGTATCTTATCATGTCTGTATACCGTCGACCTCTAGCTAGAGCTTGG
CGTAATCATGGTCATAGCTGTTTCCTGTGTGAAATTGTTATCCGCTCACAATTCCACACAACAT
ACGAGCCGGAAGCATAAAGTGTAAAGCCTGGGGTGCCTAATGAGTGAGCTAACTCACATTAAT
TGCGTTGCGCTCACTGCCCGCTTTCCAGTCGGGAAACCTGTCGTGCCAGCTGCATTAATGAAT
CGGCCAACGCGCGGGGAGAGGCGGTTTGCGTATTGGGCGCTCTTCCGCTTCCTCGCTCACTG
ACTCGCTGCGCTCGGTCGTTCGGCTGCGGCGAGCGGTATCAGCTCACTCAAAGGCGGTAATAC
GGTTATCCACAGAATCAGGGGATAACGCAGGAAAGAACATGTGAGCAAAAGGCCAGCAAAA
GGCCAGGAACCGTAAAAAGGCCGCGTTGCTGGCGTTTTTCCATAGGCTCCGCCCCCCTGACG
AGCATCACAAAAATCGACGCTCAAGTCAGAGGTGGCGAAACCCGACAGGACTATAAAGATAC
CAGGCGTTTCCCCCTGGAAGCTCCCTCGTGCGCTCTCCTGTTCCGACCCTGCCGCTTACCGGA
TACCTGTCCGCCTTTCTCCCTTCGGGAAGCGTGGCGCTTTCTCATAGCTCACGCTGTAGGTATC
TCAGTTCGGTGTAGGTCGTTCGCTCCAAGCTGGGCTGTGTGCACGAACCCCCCGTTCAGCCC
GACCGCTGCGCCTTATCCGGTAACTATCGTCTTGAGTCCAACCCGGTAAGACACGACTTATCG
CCACTGGCAGCAGCCACTGGTAACAGGATTAGCAGAGCGAGGTATGTAGGCGGTGCTACAGA
GTTCTTGAAGTGGTGGCCTAACTACGGCTACACTAGAAGAACAGTATTTGGTATCTGCGCTCT
GCTGAAGCCAGTTACCTTCGGAAAAAGAGTTGGTAGCTCTTGATCCGGCAAACAAACCACCG
CTGGTAGCGGTGGTTTTTTTGTTTGCAAGCAGCAGATTACGCGCAGAAAAAAAGGATCTCAA
GAAGATCCTTTGATCTTTTCTACGGGGTCTGACGCTCAGTGGAACGAAAACTCACGTTAAGGG
ATTTTGGTCATGAGATTATCAAAAAGGATCTTCACCTAGATCCTTTTAAATTAAAAATGAAGTT
TTAAATCAATCTAAAGTATATATGAGTAAACTTGGTCTGACAGTTACCAATGCTTAATCAGTGA
GGCACCTATCTCAGCGATCTGTCTATTTCGTTCATCCATAGTTGCCTGACTCCCCGTCGTGTAG
ATAACTACGATACGGGAGGGCTTACCATCTGGCCCCAGTGCTGCAATGATACCGCGAGACCCA
CGCTCACCGGCTCCAGATTTATCAGCAATAAACCAGCCAGCCGGAAGGGCCGAGCGCAGAAG
TGGTCCTGCAACTTTATCCGCCTCCATCCAGTCTATTAATTGTTGCCGGGAAGCTAGAGTAAGT
AGTTCGCCAGTTAATAGTTTGCGCAACGTTGTTGCCATTGCTACAGGCATCGTGGTGTCACGC
TCGTCGTTTGGTATGGCTTCATTCAGCTCCGGTTCCCAACGATCAAGGCGAGTTACATGATCCC
CCATGTTGTGCAAAAAAGCGGTTAGCTCCTTCGGTCCTCCGATCGTTGTCAGAAGTAAGTTGG
CCGCAGTGTTATCACTCATGGTTATGGCAGCACTGCATAATTCTCTTACTGTCATGCCATCCGTA
AGATGCTTTTCTGTGACTGGTGAGTACTCAACCAAGTCATTCTGAGAATAGTGTATGCGGCGA
CCGAGTTGCTCTTGCCCGGCGTCAATACGGGATAATACCGCGCCACATAGCAGAACTTTAAAA
GTGCTCATCATTGGAAAACGTTCTTCGGGGCGAAAACTCTCAAGGATCTTACCGCTGTTGAGA
TCCAGTTCGATGTAACCCACTCGTGCACCCAACTGATCTTCAGCATCTTTTACTTTCACCAGCG
TTTCTGGGTGAGCAAAAACAGGAAGGCAAAATGCCGCAAAAAAGGGAATAAGGGCGACACG
GAAATGTTGAATACTCATACTCTTCCTTTTTCAATATTATTGAAGCATTTATCAGGGTTATTGTCT
CATGAGCGGATACATATTTGAATGTATTTAGAAAAATAAACAAATAGGGGTTCCGCGCACATTT
CCCCGAAAAGTGCCACCTGACGTC

b. **Figure S2: Fusion Plasmid Map**

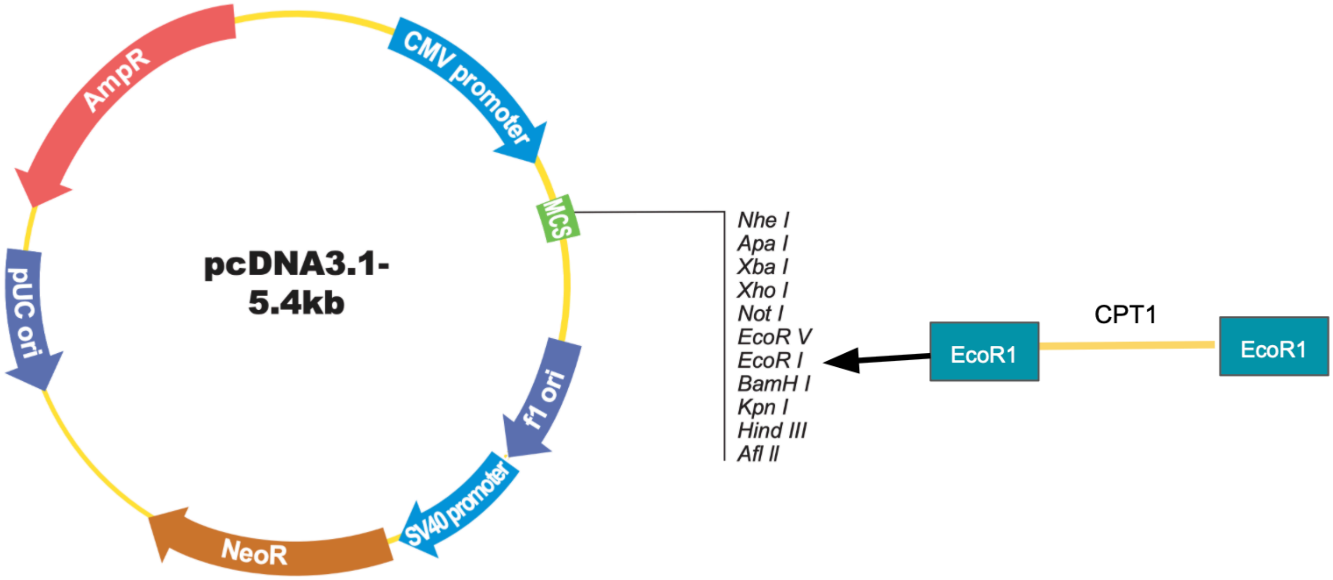

### C. CPT1B Non-fusion Plasmid

#### a. Non-fusion Plasmid Sequence

pcDNA3.1
> CPT1B(NM) ORF Clone
GACGGATCGGGAGATCTCCCGATCCCCTATGGTGCACTCTCAGTACAATCTGCTCTGATGCCG
CATAGTTAAGCCAGTATCTGCTCCCTGCTTGTGTGTTGGAGGTCGCTGAGTAGTGCGCGAGCA
AAATTTAAGCTACAACAAGGCAAGGCTTGACCGACAATTGCATGAAGAATCTGCTTAGGGTTA
GGCGTTTTGCGCTGCTTCGCGATGTACGGGCCAGATATACGCGTTGACATTGATTATTGACTAG
TTATTAATAGTAATCAATTACGGGGTCATTAGTTCATAGCCCATATATGGAGTTCCGCGTTACATA
ACTTACGGTAAATGGCCCGCCTGGCTGACCGCCCAACGACCCCCGCCCATTGACGTCAATAAT
GACGTATGTTCCCATAGTAACGCCAATAGGGACTTTCCATTGACGTCAATGGGTGGAGTATTTA
CGGTAAACTGCCCACTTGGCAGTACATCAAGTGTATCATATGCCAAGTACGCCCCCTATTGACG
TCAATGACGGTAAATGGCCCGCCTGGCATTATGCCCAGTACATGACCTTATGGGACTTTCCTAC
TTGGCAGTACATCTACGTATTAGTCATCGCTATTACCATGGTGATGCGGTTTTGGCAGTACATCA
ATGGGCGTGGATAGCGGTTTGACTCACGGGGATTTCCAAGTCTCCACCCCATTGACGTCAATG
GGAGTTTGTTTTGGCACCAAAATCAACGGGACTTTCCAAAATGTCGTAACAACTCCGCCCCAT
TGACGCAAATGGGCGGTAGGCGTGTACGGTGGGAGGTCTATATAAGCAGAGCTCTCTGGCTAA
CTAGAGAACCCACTGCTTACTGGCTTATCGAAATTAATACGACTCACTATAGGGAGACCCAAG
CTGGCTAGCGTTTAAACTTAAGCTTGGTACCGAGCTCGGATCCGCCACCATGGCTGAGGCCCA
CCAGGCCGTGGCCTTCCAGTTCACTGTGACACCCGACGGCGTGGACTTCCGGCTGTCTAGAG
AAGCTCTGAAGCACGTGTATCTGAGCGGCATCAACAGCTGGAAGAAGAGACTGATCAGAATC
AAGAACGGCATTCTGAGGGGTGTGTACCCCGGCAGCCCTACCAGCTGGCTGGTGGTCATCAT
GGCCACAGTGGGATCATCTTTCTGCAACGTGGACATCAGCCTGGGCCTGGTCAGCTGTATCCA
GCGGTGCCTGCCTCAGGGCTGCGGCCCTTACCAAACACCTCAGACAAGAGCCCTGCTGAGCA
TGGCCATCTTCAGCACCGGCGTGTGGGTGACCGGCATCTTCTTCTTTCGGCAGACCCTGAAGC
TGCTGCTGTGCTACCACGGATGGATGTTCGAGATGCACGGCAAAACCTCCAACCTGACCAGA
ATCTGGGCTATGTGCATCCGCCTGCTGAGCAGCAGACACCCTATGCTGTATAGCTTCCAGACC
AGCCTGCCTAAGCTGCCTGTGCCTAGAGTGTCCGCCACCATCCAGAGATACCTGGAAAGCGT
GCGGCCTCTGCTCGACGATGAGGAATACTACCGGATGGAACTGCTGGCCAAAGAGTTCCAAG
ATAAGACCGCCCCTCGGCTGCAGAAATACCTGGTGCTGAAGAGCTGGTGGGCCAGCAACTAC
GTGTCTGATTGGTGGGAGGAATACATCTACCTGAGAGGCAGATCCCCACTGATGGTGAACTCT
AATTACTACGTGATGGACCTGGTGCTGATCAAGAATACCGACGTGCAGGCCGCTCGTCTGGGC
AACATCATCCACGCCATGATCATGTACCGGAGAAAGCTGGATAGAGAAGAAATCAAACCTGTG
ATGGCCCTTGGCATCGTGCCCATGTGTAGCTACCAGATGGAAAGAATGTTCAACACCACCAGG
ATCCCTGGCAAGGACACCGATGTTCTGCAGCATCTCAGCGACAGCAGACACGTTGCCGTGTA
CCACAAGGGCCGGTTCTTTAAGCTGTGGCTGTACGAGGGCGCCAGACTGCTGAAACCTCAGG
ACCTGGAAATGCAGTTCCAGCGGATCCTGGACGACCCGAGCCCCCCCAACCTGGAGAAGAG
AAGCTGGCTGCCCTGACCGCCGGCGGAAGAGTGGAATGGGCTCAGGCCAGACAGGCATTCTT
CAGCTCTGGCAAGAACAAGGCCGCCCTGGAAGCCATCGAGAGAGCCGCTTTTTTCGTGGCTC
TCGACGAGGAAAGCTACAGCTACGACCCCGAGGACGAGGCCTCTCTGAGCCTGTACGGCAA
GGCCCTGCTGCACGGCAACTGCTACAACAGATGGTTTGATAAGTCCTTCACCCTGATCAGCTT
CAAGAATGGCCAGCTGGGACTGAACGCCGAGCACGCCTGGGCCGACGCCCCTATCATCGGCC
ACCTGTGGGAATTCGTGCTAGGTACAGATAGCTTCCACCTGGGATATACAGAGACAGGCCACT

GCCTGGGCAAGCCCAACCCCGCCCTGGCCCCTCCTACAAGACTGCAGTGGGACATCCCTAAG
CAGTGTCAGGCAGTGATCGAGAGCTCTTATCAAGTGGCCAAAGCTCTGGCCGATGACGTGGA
GCTGTACTGCTTCCAGTTTCTGCCATTCGGCAAGGGCCTGATCAAGAAGTGCAGAACCAGCC
CAGACGCCTTCGTGCAGATCGCCCTCCAGCTGGCGCATTTCAGAGATAGAGGCAAATTCTGTT
TAACATACGAGGCCAGCATGACCAGAATGTTCAGAGAGGGCCGGACCGAGACCGTGCGGAG
CTGCACCAGCGAGTCTACAGCCTTCGTACAGGCCATGATGGAAGGATCTCACACCAAGGCCG
ATCTGCGGGATCTCTTCCAGAAGGCTGCTAAGAAACACCAGAACATGTACAGACTCGCCATG
ACAGGGGCCGGCATCGACAGACATCTGTTTGCCTGTACCTGGTGTCCAAGTACCTGGGTGTG
TCCAGCCCCTTCCTGGCCGAGGTGCTGTCCGAGCCTTGGAGACTGTCTACCAGCCAGATTCCT
CAAAGCCAAATCAGAATGTTCGATCCTGAGCAACACCCCAATCACCTGGGCGCCGGAGGCGG
ATTTGGCCCCGTGGCCGACGACGGCTACGGCGTCAGTTATATGATCGCAGGCGAGAACACCAT
CTTTTTTCACATCAGTTCCAAGTTCTCATCTAGCGAGACAAACGCTCAGAGATTCGGCAATCA
CATTCGAAAAGCTCTGCTGGACATCGCCGACCTGTTCCAAGTGCCAAAGGCCTACAGCGATTA
CAAGGATGACGACGATAAGTGATAAACCCGCTGATCAGCCTCGACTGTGCCTTCTAGTTGCCA
GCCATCTGTTGTTTGCCCCTCCCCCGTGCCTTCCTTGACCCTGGAAGGTGCCACTCCCACTGT
CCTTTCCTAATAAAATGAGGAAATTGCATCGCATTGTCTGAGTAGGTGTCATTCTATTCTGGGG
GGTGGGGTGGGGCAGGACAGCAAGGGGGAGGATTGGGAAGACAATAGCAGGCATGCTGGGG
ATGCGGTGGGCTCTATGGCTTCTGAGGCGGAAAGAACCAGCTGGGGCTCTAGGGGGTATCCC
CACGCGCCCTGTAGCGGCGCATTAAGCGCGGCGGGTGTGGTGGTTACGCGCAGCGTGACCGC
TACACTTGCCAGCGCCCTAGCGCCCGCTCCTTTCGCTTTCTTCCCTTCCTTTCTCGCCACGTTC
GCCGGCTTTCCCCGTCAAGCTCTAAATCGGGGGCTCCCTTTAGGGTTCCGATTTAGTGCTTTAC
GGCACCTCGACCCCAAAAAACTTGATTAGGGTGATGGTTCACGTAGTGGGCCATCGCCCTGAT
AGACGGTTTTTCGCCCTTTGACGTTGGAGTCCACGTTCTTTAATAGTGGACTCTTGTTCCAAAC
TGGAACAACACTCAACCCTATCTCGGTCTATTCTTTTGATTTATAAGGGATTTTGCCGATTTCGG
CCTATTGGTTAAAAAATGAGCTGATTTAACAAAAATTTAACGCGAATTAATTCTGTGGAATGTG
TGTCAGTTAGGGTGTGGAAAGTCCCCAGGCTCCCCAGCAGGCAGAAGTATGCAAAGCATGCA
TCTCAATTAGTCAGCAACCAGGTGTGGAAAGTCCCCAGGCTCCCCAGCAGGCAGAAGTATGC
AAAGCATGCATCTCAATTAGTCAGCAACCATAGTCCCGCCCCTAACTCCGCCCATCCCGCCCCT
AACTCCGCCCAGTTCCGCCCATTCTCCGCCCCATGGCTGACTAATTTTTTTTATTTATGCAGAG
GCCGAGGCCGCCTCTGCCTCTGAGCTATTCCAGAAGTAGTGAGGAGGCTTTTTTGGAGGCCTA
GGCTTTTGCAAAAAGCTCCCGGGAGCTTGTATATCCATTTTCGGATCTGATCAAGAGACAGGA
TGAGGATCGTTTCGCATGATTGAACAAGATGGATTGCACGCAGGTTCTCCGGCCGCTTGGGTG
GAGAGGCTATTCGGCTATGACTGGGCACAACAGACAATCGGCTGCTCTGATGCCGCCGTGTTC
CGGCTGTCAGCGCAGGGGCGCCCGGTTCTTTTTGTCAAGACCGACCTGTCCGGTGCCCTGAA
TGAACTGCAGGACGAGGCAGCGCGGCTATCGTGGCTGGCCACGACGGGCGTTCCTTGCGCAG
CTGTGCTCGACGTTGTCACTGAAGCGGGAAGGGACTGGCTGCTATTGGGCGAAGTGCCGGGG
CAGGATCTCCTGTCATCTCACCTTGCTCCTGCCGAGAAAGTATCCATCATGGCTGATGCAATGC
GGCGGCTGCATACGCTTGATCCGGCTACCTGCCCATTCGACCACCAAGCGAAACATCGCATCG
AGCGAGCACGTACTCGGATGGAAGCCGGTCTTGTCGATCAGGATGATCTGGACGAAGAGCAT
CAGGGGCTCGCGCCAGCCGAACTGTTCGCCAGGCTCAAGGCGCGCATGCCCGACGGCGAGG
ATCTCGTCGTGACCCATGGCGATGCCTGCTTGCCGAATATCATGGTGGAAAATGGCCGCTTTTC
TGGATTCATCGACTGTGGCCGGCTGGGTGTGGCGGACCGCTATCAGGACATAGCGTTGGCTAC
CCGTGATATTGCTGAAGAGCTTGGCGGCGAATGGGCTGACCGCTTCCTCGTGCTTTACGGTAT
CGCCGCTCCCGATTCGCAGCGCATCGCCTTCTATCGCCTTCTTGACGAGTTCTTCTGAGCGGG

ACTCTGGGGTTCGAAATGACCGACCAAGCGACGCCCAACCTGCCATCACGAGATTTCGATTCC
ACCGCCGCCTTCTATGAAAGGTTGGGCTTCGGAATCGTTTTCCGGGACGCCGGCTGGATGATC
CTCCAGCGCGGGGATCTCATGCTGGAGTTCTTCGCCCACCCCAACTTGTTTATTGCAGCTTATA
ATGGTTACAAATAAAGCAATAGCATCACAAATTTCACAAATAAAGCATTTTTTTCACTGCATTC
TAGTTGTGGTTTGTCCAAACTCATCAATGTATCTTATCATGTCTGTATACCGTCGACCTCTAGCT
AGAGCTTGGCGTAATCATGGTCATAGCTGTTTCCTGTGTGAAATTGTTATCCGCTCACAATTCC
ACACAACATACGAGCCGGAAGCATAAAGTGTAAAGCCTGGGGTGCCTAATGAGTGAGCTAAC
TCACATTAATTGCGTTGCGCTCACTGCCCGCTTTCCAGTCGGGAAACCTGTCGTGCCAGCTGC
ATTAATGAATCGGCCAACGCGCGGGGAGAGGCGGTTTGCGTATTGGGCGCTCTTCCGCTTCCT
CGCTCACTGACTCGCTGCGCTCGGTCGTTCGGCTGCGGCGAGCGGTATCAGCTCACTCAAAG
GCGGTAATACGGTTATCCACAGAATCAGGGGATAACGCAGGAAAGAACATGTGAGCAAAAGG
CCAGCAAAAGGCCAGGAACCGTAAAAAGGCCGCGTTGCTGGCGTTTTTCCATAGGCTCCGCC
CCCCTGACGAGCATCACAAAAATCGACGCTCAAGTCAGAGGTGGCGAAACCCGACAGGACT
ATAAAGATACCAGGCGTTTCCCCCTGGAAGCTCCCTCGTGCGCTCTCCTGTTCCGACCCTGCC
GCTTACCGGATACCTGTCCGCCTTTCTCCCTTCGGGAAGCGTGGCGCTTTCTCATAGCTCACGC
TGTAGGTATCTCAGTTCGGTGTAGGTCGTTCGCTCCAAGCTGGGCTGTGTGCACGAACCCCCC
GTTCAGCCCGACCGCTGCGCCTTATCCGGTAACTATCGTCTTGAGTCCAACCCGGTAAGACAC
GACTTATCGCCACTGGCAGCAGCCACTGGTAACAGGATTAGCAGAGCGAGGTATGTAGGCGG
TGCTACAGAGTTCTTGAAGTGGTGGCCTAACTACGGCTACACTAGAAGAACAGTATTTGGTAT
CTGCGCTCTGCTGAAGCCAGTTACCTTCGGAAAAAGAGTTGGTAGCTCTTGATCCGGCAAAC
AAACCACCGCTGGTAGCGGTGGTTTTTTTGTTTGCAAGCAGCAGATTACGCGCAGAAAAAAA
GGATCTCAAGAAGATCCTTTGATCTTTTCTACGGGGTCTGACGCTCAGTGGAACGAAAACTCA
CGTTAAGGGATTTTGGTCATGAGATTATCAAAAAGGATCTTCACCTAGATCCTTTTAAATTAAA
AATGAAGTTTTAAATCAATCTAAAGTATATATGAGTAAACTTGGTCTGACAGTTACCAATGCTT
AATCAGTGAGGCACCTATCTCAGCGATCTGTCTATTTCGTTCATCCATAGTTGCCTGACTCCCC
GTCGTGTAGATAACTACGATACGGGAGGGCTTACCATCTGGCCCCAGTGCTGCAATGATACCG
CGAGACCCACGCTCACCGGCTCCAGATTTATCAGCAATAAACCAGCCAGCCGGAAGGGCCGA
GCGCAGAAGTGGTCCTGCAACTTTATCCGCCTCCATCCAGTCTATTAATTGTTGCCGGGAAGCT
AGAGTAAGTAGTTCGCCAGTTAATAGTTTGCGCAACGTTGTTGCCATTGCTACAGGCATCGTG
GTGTCACGCTCGTCGTTTGGTATGGCTTCATTCAGCTCCGGTTCCCAACGATCAAGGCGAGTT
ACATGATCCCCCATGTTGTGCAAAAAAGCGGTTAGCTCCTTCGGTCCTCCGATCGTTGTCAGA
AGTAAGTTGGCCGCAGTGTTATCACTCATGGTTATGGCAGCACTGCATAATTCTCTTACTGTCA
TGCCATCCGTAAGATGCTTTTCTGTGACTGGTGAGTACTCAACCAAGTCATTCTGAGAATAGT
GTATGCGGCGACCGAGTTGCTCTTGCCCGGCGTCAATACGGGATAATACCGCGCCACATAGCA
GAACTTTAAAAGTGCTCATCATTGGAAAACGTTCTTCGGGGCGAAAACTCTCAAGGATCTTAC
CGCTGTTGAGATCCAGTTCGATGTAACCCACTCGTGCACCCAACTGATCTTCAGCATCTTTTAC
TTTCACCAGCGTTTCTGGGTGAGCAAAAACAGGAAGGCAAAATGCCGCAAAAAAGGGAATA
AGGGCGACACGGAAATGTTGAATACTCATACTCTTCCTTTTTCAATATTATTGAAGCATTTATCA
GGGTTATTGTCTCATGAGCGGATACATATTTGAATGTATTTAGAAAAATAAACAAATAGGGGTT
CCGCGCACATTTCCCCGAAAAGTGCCACCTGACGTC

**b. Figure S3: Non-fusion Plasmid Map**

**After the insertion following common subclone protocols, we inserted our CPT1b wild-type gene into our pcDNA 3.1 vector into the cloning MCS site. After we sequenced, there is no new mutation, we validated the start point and the stop codon works well. There is no reading frameshift mutation. We then used this vector for our CPT1a and CPT1b protein harvest.**

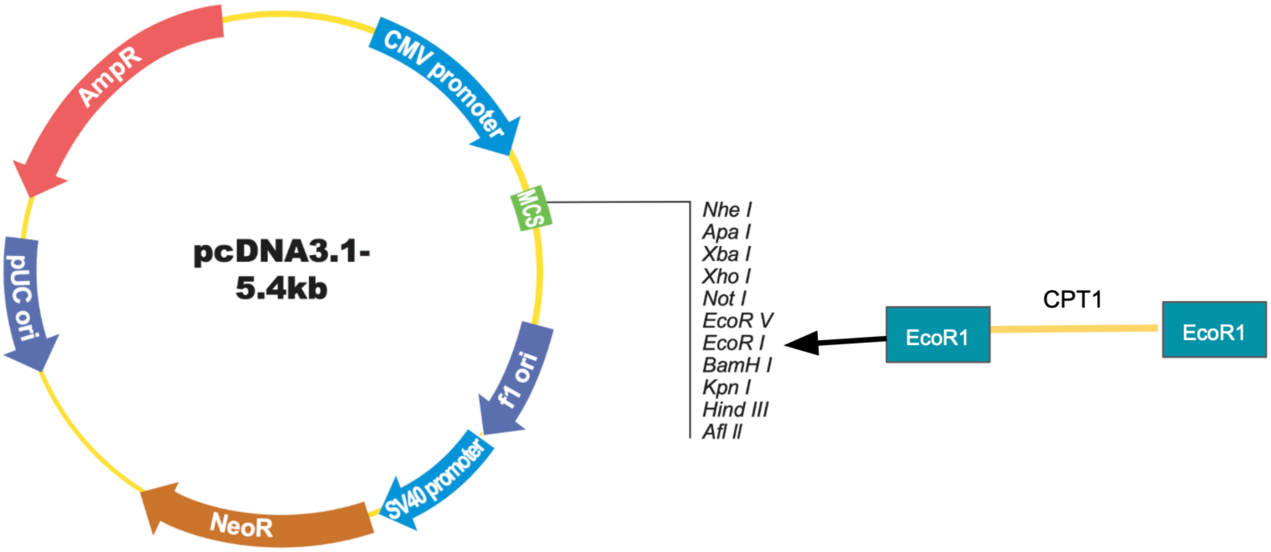

### D. CPT1B Fusion Plasmid

#### a. Fusion Plasmid Sequence

>CPT1B(NM_897968) ORF Clone
ATGGCTGAGGCCCACCAGGCCGTGGCCTTCCAGTTCACTGTGACACCCGACGGCGTGGACTT
CCGGCTGTCTAGAGAAGCTCTGAAGCACGTGTATCTGAGCGGCATCAACAGCTG
GAAGAAGAGACTGATCAGAATCAAGAACGGCATTCTGAGGGGTGTGTACCCCGGCAGCCCTA
CCAGCTGGCTGGTGGTCATCATGGCCACAGTGGGATCATCTTTCTGCAACGTGG
ACATCAGCCTGGGCCTGGTCAGCTGTATCCAGCGGTGCCTGCCTCAGGGCTGCGGCCCTTACC
AAACACCTCAGACAAGAGCCCTGCTGAGCATGGCCATCTTCAGCACCGGCGTG
TGGGTGACCGGCATCTTCTTCTTTCGGCAGACCCTGAAGCTGCTGCTGTGCTACCACGGATGG
ATGTTCGAGATGCACGGCAAAACCTCCAACCTGACCAGAATCTGGGCTATGTGC
ATCCGCCTGCTGAGCAGCAGACACCCTATGCTGTATAGCTTCCAGACCAGCCTGCCTAAGCTG
CCTGTGCCTAGAGTGTCCGCCACCATCCAGAGATACCTGGAAAGCGTGCGGCCT
CTGCTCGACGATGAGGAATACTACCGGATGGAACTGCTGGCCAAAGAGTTCCAAGATAAGAC
CGCCCCTCGGCTGCAGAAATACCTGGTGCTGAAGAGCTGGTGGGCCAGCAACTA
CGTGTCTGATTGGTGGGAGGAATACATCTACCTGAGAGGCAGATCCCCACTGATGGTGAACTC
TAATTACTACGTGATGGACCTGGTGCTGATCAAGAATACCGACGTGCAGGCCGC
TCGTCTGGGCAACATCATCCACGCCATGATCATGTACCGGAGAAAGCTGGATAGAGAAGAAAT
CAAACCTGTGATGGCCCTTGGCATCGTGCCCATGTGTAGCTACCAGATGGAAAG
AATGTTCAACACCACCAGGATCCCTGGCAAGGACACCGATGTTCTGCAGCATCTCAGCGACA
GCAGACACGTTGCCGTGTACCACAAGGGCCGGTTCTTTAAGCTGTGGCTGTACGA
GGGCGCCAGACTGCTGAAACCTCAGGACCTGGAAATGCAGTTCCAGCGGATCCTGGACGACC
CGAGCCCCCCCAACCTGGAGAAGAGAAGCTGGCTGCCCTGACCGCCGGCGGA
AGAGTGGAATGGGCTCAGGCCAGACAGGCATTCTTCAGCTCTGGCAAGAACAAGGCCGCCCT
GGAAGCCATCGAGAGAGCCGCTTTTTCGTGGCTCTCGACGAGGAAAGCTACAG
CTACGACCCCGAGGACGAGGCCTCTCTGAGCCTGTACGGCAAGGCCCTGCTGCACGGCAACT
GCTACAACAGATGGTTTGATAAGTCCTTCACCCTGATCAGCTTCAAGAATGGCCA
GCTGGGACTGAACGCCGAGCACGCCTGGGCCGACGCCCCTATCATCGGCCACCTGTGGGAAT
TCGTGCTAGGTACAGATAGCTTCCACCTGGGATATACAGAGACAGGCCACTGCC
TGGGCAAGCCCAACCCCGCCCTGGCCCCTCCTACAAGACTGCAGTGGGACATCCCTAAGCAG
TGTCAGGCAGTGATCGAGAGCTCTTATCAAGTGGCCAAAGCTCTGGCCGATGAC
GTGGAGCTGTACTGCTTCCAGTTTCTGCCATTCGGCAAGGGCCTGATCAAGAAGTGCAGAAC
CAGCCCAGACGCCTTCGTGCAGATCGCCCTCCAGCTGGCGCATTTCAGAGATAGA
GGCAAATTCTGTTTAACATACGAGGCCAGCATGACCAGAATGTTCAGAGAGGGCCGGACCGA
GACCGTGCGGAGCTGCACCAGCGAGTCTACAGCCTTCGTACAGGCCATGATGGA
AGGATCTCACACCAAGGCCGATCTGCGGGATCTCTTCCAGAAGGCTGCTAAGAAACACCAGA
ACATGTACAGACTCGCCATGACAGGGGCCGGCATCGACAGACATCTGTTTTGCCT
GTACCTGGTGTCCAAGTACCTGGGTGTGTCCAGCCCCTTCCTGGCCGAGGTGCTGTCCGAGCC
TTGGAGACTGTCTACCAGCCAGATTCCTCAAAGCCAAATCAGAATGTTCGATCC
TGAGCAACACCCCAATCACCTGGGCGCCGGAGGCGGATTTGGCCCCGTGGCCGACGACGGCT
ACGGCGTCAGTTATATGATCGCAGGCGAGAACACCATCTTTTTCACATCAGTTC

CAAGTTCTCATCTAGCGAGACAAACGCTCAGAGATTCGGCAATCACATTCGAAAAGCTCTGCT
GGACATCGCCGACCTGTTCCAAGTGCCAAAGGCCTACAGC

#### b. Figure S4: Fusion Plasmid Map

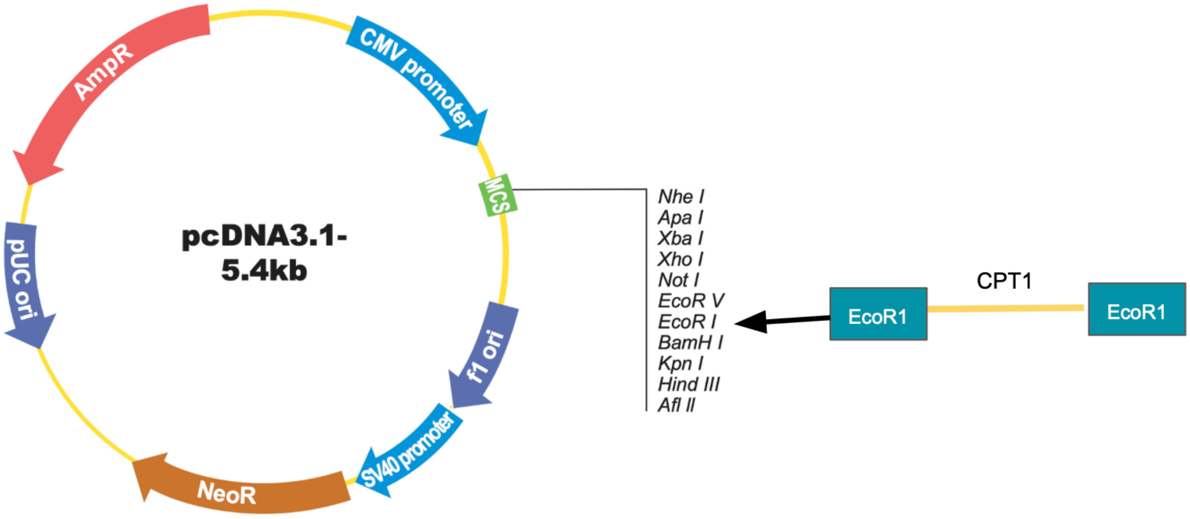

#### E. Figure S5: Standard Curve for DTNB

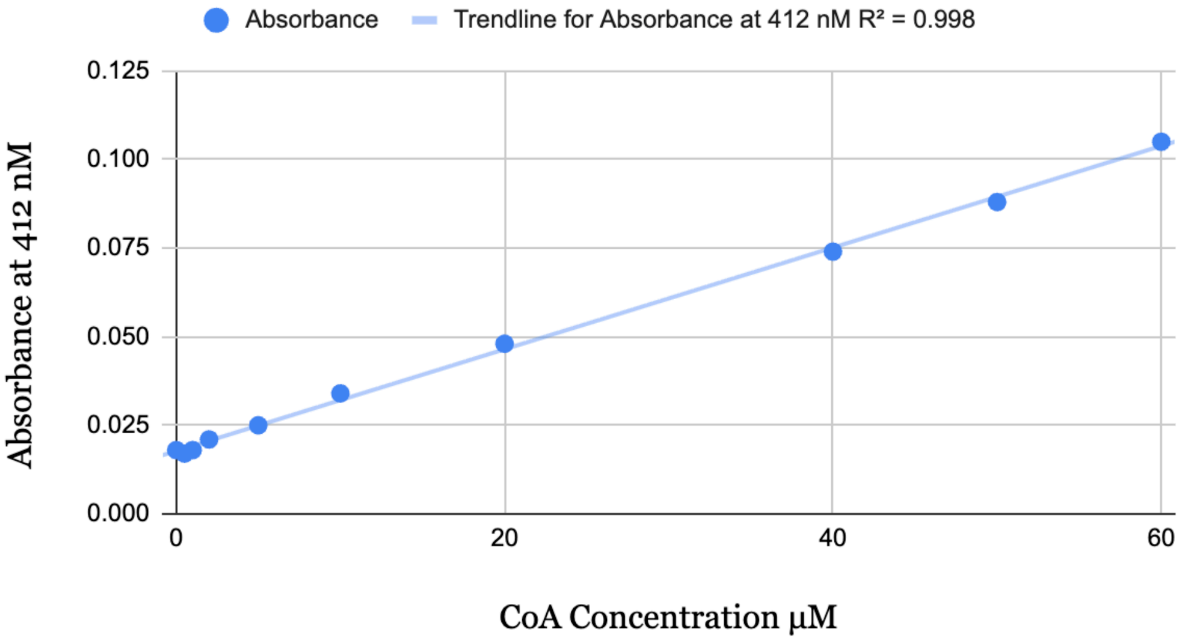


#### F. Figure S6: Standard Curve for Protein Concentration

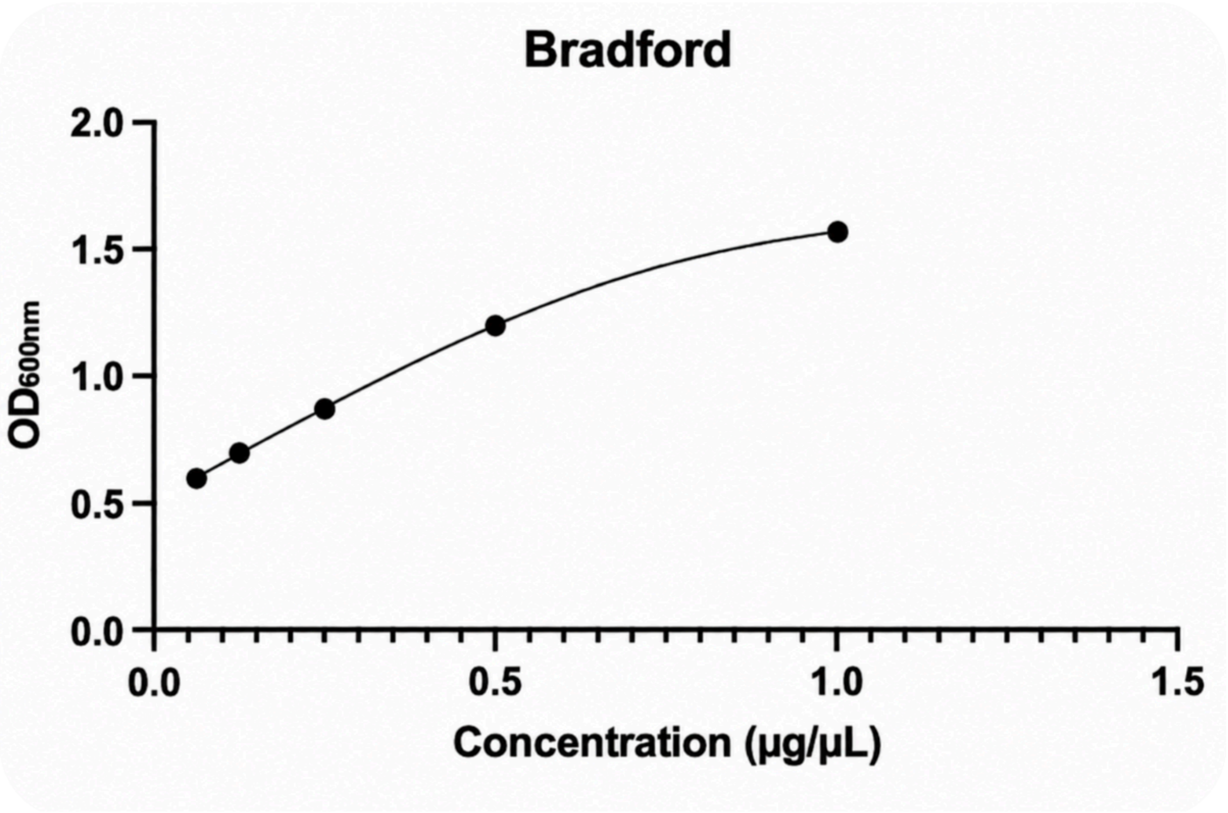
